# Humans use overconfident estimates of auditory spatial and temporal uncertainty for perceptual inference

**DOI:** 10.1101/2024.12.01.626278

**Authors:** Fangfang Hong, Jiaming Xu, Michael S. Landy, Stephanie Badde

## Abstract

Making decisions based on noisy sensory information is a crucial function of the brain. Various decisions take each sensory signal’s uncertainty into account. Here, we investigated whether perceptual inferences rely on accurate estimates of sensory uncertainty. Participants completed a set of auditory, visual, and audiovisual spatial as well as temporal tasks. We fitted Bayesian observer models of each task to every participant’s complete dataset. Crucially, in some model variants the uncertainty estimates employed for perceptual inferences were independent of the actual uncertainty associated with the sensory signals. Model comparisons and analysis of the best-fitting parameters revealed that, in unimodal and bimodal contexts, participants’ perceptual decisions relied on overconfident estimates of auditory spatial and audiovisual temporal uncertainty. These findings challenge the ubiquitous assumption that human behavior optimally accounts for sensory uncertainty regardless of sensory domain.

## Introduction

Our senses are inherently noisy [1]; repeated exposure to the same physical stimulus rarely leads to the same neural activity in the brain and, thus, rarely to the same sensory measurement of the stimulus properties (**Fig. 1A-B**). Imagine your dog breaks free from its leash while you are in the woods. You hear a series of your dog’s barks. Due to sensory noise, the sensory measurements of the barks’ locations vary even if the dog stays in place. The probability distribution of the spatial measurements across many barks (the “measurement distribution”) reflects auditory spatial uncertainty; the broader the distribution, the higher the level of uncertainty. Consequently, a single bark might have originated from a range of plausible locations, each with its own likelihood of being the true origin (**Fig. 1C**). If decisions take sensory uncertainty into account, i.e., consider all possible locations and their likelihood (the “likelihood function”), changes in sensory uncertainty should lead to specific changes in behavior. Indeed, the degree of uncertainty associated with sensory signals influences our perception [2], cognition [3], action [4, 5], memory [6], learning, and decision-making [7, 8], indicating that a wide range of human behaviors takes sensory uncertainty into account [9].

**Figure 1.**
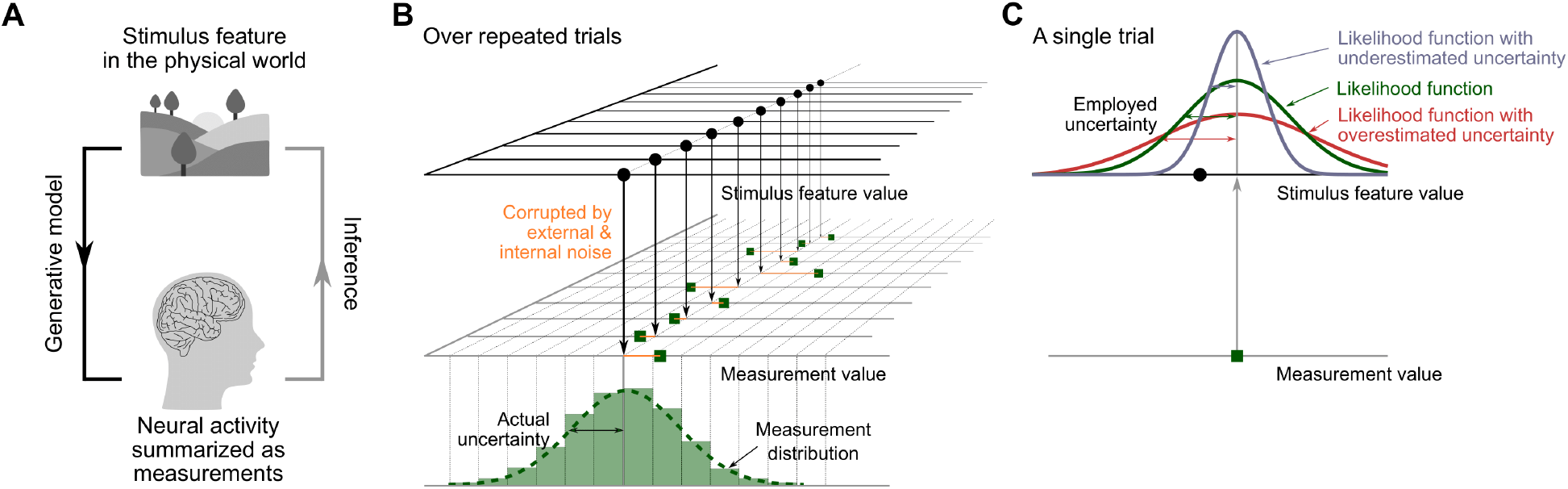
Generative model and perceptual inference. (A) An event in the world leads to neural activity that represents stimulus properties such as the stimulus location. The brain only has access to the measurement and has to infer the stimulus property that gave rise to it. (B) Due to external and internal noise, the measurements will not be aligned perfectly with the stimulus property in the physical world. The standard deviation of the measurements over repeated presentations of the same stimulus describes the observer’s actual sensory uncertainty. (C) In a single trial, the brain infers the physical property of the stimulus based on a single measurement; perceptual inference relies on the likelihood of each possible value of the stimulus property given both the measurement and its estimated uncertainty. For a symmetric measurement distribution, the shape of the likelihood function equals that of the measurement distribution if the uncertainty is estimated accurately.

The match between sensory uncertainty estimates underlying perception and actual uncertainty remains contested. Bayesian encoding theory posits that uncertainty is automatically encoded in the activity of neurons selective for the stimulus feature. According to this account, the likelihood can be inferred from a probabilistic population code [10-13], implying that uncertainty is accurately represented and employed to mediate various decisions. This theory is supported by neurophysiological evidence showing that the likelihood function can be decoded from population activity in primary visual cortex. Monkey behavior in an orientation-classification task is better predicted by models using trial-specific estimates of uncertainty from neural responses, compared to a model that only derives a point estimate of orientation, suggesting that trial-specific likelihood functions underlie perceptual decisions [14]. In humans, likelihood functions have been decoded from fMRI measurements of visual cortical activity and linked to performance in visual orientation estimation [15] and spatial working memory [16] tasks. Few psychophysical studies provide direct evidence that internal likelihood functions accurately mirror sensory uncertainty (**Fig. 1B,C**). These studies contrast the models with accurate or inaccurate likelihood functions in a visual spatial-localization task [17] and a visual orientation change-detection task [18]. They conclude that uncertainty is automatically and accurately encoded as a likelihood function. These studies focus on visual location or orientation, known to be encoded by tuned populations of visual neurons. Other studies directly probed the accuracy of motor uncertainty estimates employed in visuomotor tasks. They revealed misestimation of motor uncertainty, finding either under-or overestimation of uncertainty [19, 20]. Uncertainty-based inference underpins many human behaviors, yet clear evidence that these rely on accurate uncertainty estimates has been limited to a few visual features. This lack of behavioral evidence may be due to nontrivial methodological and computational challenges.

First, many parameters influence human behavior similarly, making it challenging to attribute changes in behavior to a single parameter. Specifically, it is difficult to isolate the estimate of sensory uncertainty from Bayesian priors and biases. Let’s continue with the dog-search example. Previous experience might tell you that your dog often dashes toward its favorite rabbit hole, left of where you heard the bark. Optimally, this knowledge would steer you to the left of the bark, with the exact deviation depending on the spatial uncertainty of the bark and the precision of your spatial representation of the rabbit hole. If you nevertheless sprint straight towards the bark, there are at least three explanations for your behavior. 1) The bark provided precise spatial information, leading you to rely mostly on the auditory measurement. 2) The bark provided uncertain spatial information, but your inference relied on an overconfident estimate of that spatial uncertainty, again leading to dominance of the auditory measurement. 3) Your auditory spatial perception has a rightward bias; consequently, even though you integrated the rabbit hole prior, the final location estimate happened to align with the true location of the bark. Many studies conclude that the brain employs accurate estimates of sensory uncertainty based on good matches between human behavior and predictions of a Bayesian ideal-observer model [4, 21-24]. However, in these models, sensory uncertainty and priors trade off. Even if the prior is learned during the experiment, it might be learned imprecisely [25-27], and its final representation is inaccessible to the experimenter, as are estimates of sensory uncertainty employed for perceptual inference.

Second, the model has to capture the complexity of inferences about the world that underlie observed behavior. Continuing with the search for your dog, suppose in addition to the sound, you spot movement behind a nearby bush. Optimally combining the two pieces of information based on their relative precision will reduce spatial uncertainty [28-30]. You might aim more towards the shaking bush as vision usually provides more precise spatial information than audition [31-34]. Hence, multisensory integration provides a strong test case for uncertainty-based perceptual inference. Some studies report variance reduction predicted by optimal cue integration [21, 28-30, 35-40], while others fail to find this [41-48]. However, multisensory integration only makes sense when the visual and auditory information originate from the same event [49, 50]. If the moving twigs are far from the apparent location of the dog’s bark or you saw the movement long after hearing the sound, then you would probably sprint straight towards the bark **(Fig. 2A**). Yet, you might also steer towards the bark because you underestimated its spatial uncertainty. Both plausible scenarios, doubts about the shared origin of the sensory signals and employing inaccurate estimates of sensory uncertainty, will lead to sub-optimal cue combination. Thus, models must infer the causal structure of the sensory signals [27, 32, 49, 51-60] to draw conclusions about the estimates of sensory uncertainty underlying perceptual inference (**Fig. 2B**). Sensory biases affect causal inference [61] and priors over stimulus properties influence integration, stressing the need to account for potential tradeoffs between model parameters when accounting for the complexity of the observer’s world model.

**Figure 2.**
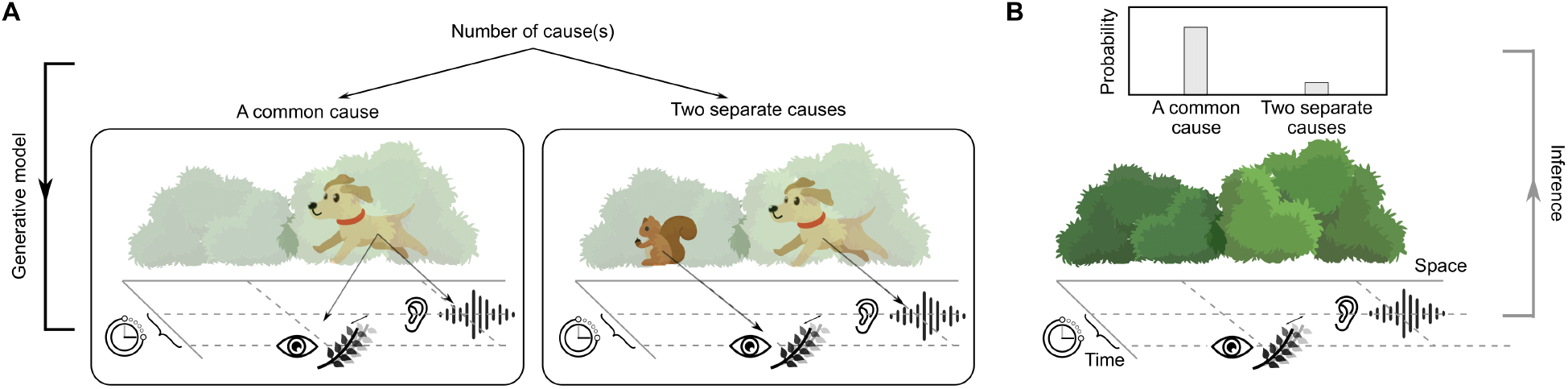
Role of causal inference in multisensory tasks. (A) A generative model describing the statistical structure of the physical world resulting in visual and auditory signals arriving in the observer’s brain. The two spatially and temporally discrepant measurements could be caused by a single source (a barking dog whose shuffling visibly shakes branches of the bush), or two different sources (a squirrel shaking the bush and a barking dog). (B) Inference about whether there is one common or two separate sources based on the visual and auditory measurements.

Here, we test whether accurate estimates of auditory temporal and spatial uncertainty underly perception across unimodal and bimodal contexts. Participants completed a series of psychophysical experiments, encompassing temporal-order judgments, spatial discrimination and localization in unimodal and bimodal contexts as well as explicit causal-inference judgments. To address the challenge of tradeoffs between parameters, we jointly fit a participant’s data from all experiments, allowing us to constrain all parameters that could explain the observed behavior. We contrasted models in which inaccurate estimates of temporal and/or spatial uncertainty were employed for perceptual inference with a model that assumed accurate uncertainty estimates. Participants typically relied on inaccurate estimates of auditory spatial uncertainty in both unimodal and bimodal contexts as well as on inaccurate estimates of audiovisual temporal uncertainty. Parameter estimates indicate that participants used overconfident estimates of auditory temporal and spatial uncertainty.

## Methods

### Participants

Thirteen participants (seven females, aged 21-34 years; twelve naive with respect to the purposes of the experiment; all right-handed) were recruited. One participant’s uncertainty estimates in the audiovisual context were excluded from one of the statistical tests, as the estimate pinned at the generously set boundary. All of them stated having no visual, auditory, or motor impairments. The experiment was approved by the institutional review board of New York University and all participants gave written informed consent prior to beginning the study.

### Apparatus and stimuli

The experiment was conducted in a dark and semi-sound-attenuated room. Participants were seated 1 m from an acoustically transparent screen (1.36 × 1.02 m, 68 × 52 deg visual angle) with their head stabilized by a chin rest. Visual stimuli were high-contrast Gaussian blobs (SD: 3.6 deg) against a black background, projected onto the screen for 100 ms. Behind the screen, a loudspeaker was mounted on a computer-controlled sledge attached to a 1.5-meter-long linear rail, which was hung from the ceiling perpendicular to the line of sight. The auditory stimuli were 100 ms-long noise bursts (20 Hz - 20.05 kHz, 60 dB) windowed using the positive half of a sine wave with a period of 200 ms. The participant’s dominant hand rested on a rollerball mouse with two buttons. Stimulus presentation, speaker movement, and data collection were controlled by an iMac running MATLAB R2017b. Visual and auditory stimuli were presented using the Psychophysics Toolbox [62].

### Procedure

In Expt. 1, participants completed a two-alternative, forced-choice audiovisual temporal-order-judgment task (**Fig. 3A**). In each trial, a visual and an auditory stimulus were presented at the central location with varying SOA (20 levels ranging from -466.67 to 466.67 ms, negative values indicate visual-first stimulus pairs). Participants reported by button press which stimulus they had perceived first. Each SOA was presented 25 times, resulting in a total of 500 trials administered in pseudorandom order. Usually participants took an hour to complete this experiment.

**Figure 3.**
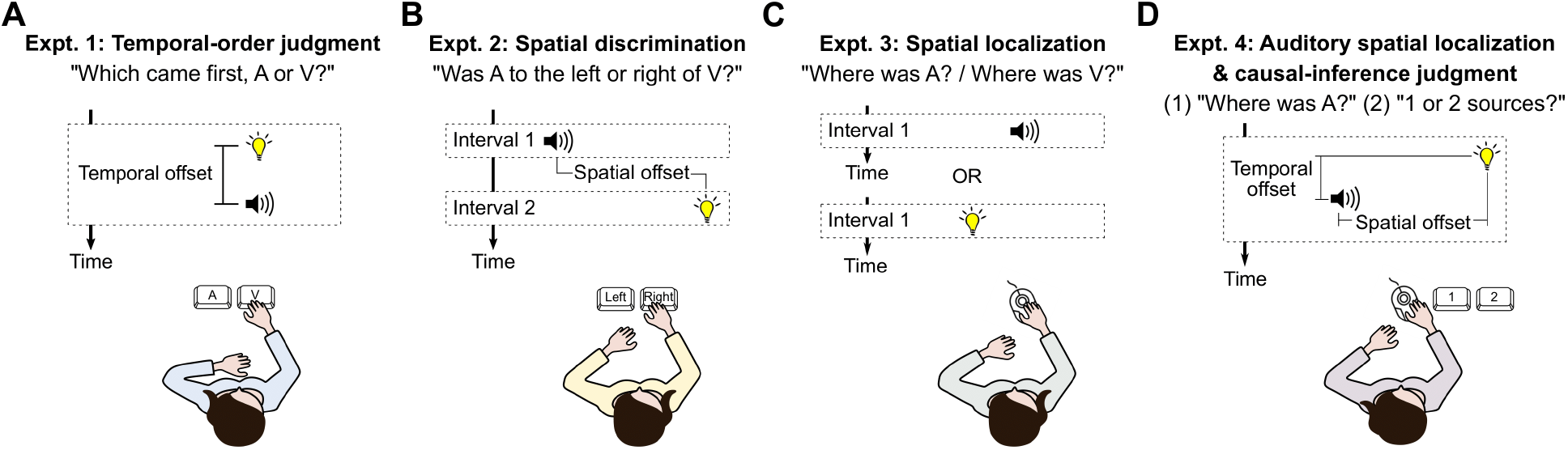
Experiments. (A) In Expt. 1, an audiovisual stimulus pair with varying temporal offset was presented at the central location. Participants reported which modality they perceived first. (B) In Expt. 2, a visual and an auditory stimulus were presented sequentially with varying spatial offset. Participants reported whether they perceived the auditory stimulus left or right of the visual stimulus. (C) In Expt. 3, either a visual or auditory stimulus was presented at a random location. Participants indicated the location of the stimulus by moving a cursor. (D) In Expt. 4, an audiovisual stimulus pair with varying spatial and temporal offsets was presented. Participants first moved a cursor to the perceived horizontal location of the auditory stimulus and then reported by button press whether they perceived the auditory and the visual stimulus as sharing a common source or originating from two separate sources.

In Expt. 2, participants completed a two-interval, forced-choice audiovisual spatial-discrimination task (**Fig. 3B**). In each trial, an auditory and a visual stimulus were presented in random order with an SOA of 1600 ms. Participants reported by button press whether they perceived the auditory stimulus to the left or right of the visual stimulus. The visual stimulus was presented at either -12 or 12 deg relative to the center of the screen. The location of the auditory stimulus was controlled by four interleaved staircases, two for each visual stimulus location. Each staircase comprised 40 trials, resulting in a total of 160 trials. Usually, participants took about an hour to complete this experiment.

In Expt. 3, participants completed an auditory and a visual spatial-localization task (**Fig. 3C**). In each trial, either an auditory or a visual stimulus was presented. Participants moved a visual cursor to the stimulus location. The visual stimulus was presented at either ±4 or ±12 deg relative to the center of the screen. Auditory stimulus locations were those for which the participant perceived auditory stimuli as aligned with visual stimuli presented at ±12 deg in Expt. 2. Each visual stimulus location was tested 20 times, and each auditory location was tested 40 times, resulting in 160 trials administered in pseudorandom order. Participants took an hour and a half to complete the experiment.

In Expt. 4, participants completed two tasks, an auditory spatial-localization task and a two-alternative, forced-choice causal-inference task (**Fig. 3D**). In each trial, an audiovisual stimulus pair was presented. Participants first moved a visual cursor to the horizontal position of the auditory stimulus, and then reported by button press whether the two stimuli originated from the same source or two separate sources. The visual stimulus was presented at ±4 or ±12 deg; the auditory stimulus was presented at one of the same two locations as in Expt. 3, those at which the auditory stimuli were perceptually aligned with visual stimuli at ±12 deg. The stimuli were presented with an SOA of 0, ±250, or ±400 ms (negative values indicate visual first). There were 40 different audiovisual stimulus pairs (4 visual stimulus locations x 2 auditory stimulus locations x 5 SOAs), each of which was tested 20 times, resulting in 800 trials. Usually, participants took three hours to complete the entire experiment.

Response feedback was never provided. The four experiments were split across three sessions. During the first session, participants completed Expts. 2 and 3; in the second session, they completed about 65% of Expt. 4; in the third session, they completed the rest of Expt. 4, followed by Expt. 1. For further experimental details, see **Supplement S1**.

## Modeling

We begin with the general assumptions of the models and then briefly describe the model variants we tested and how they were fit to the data. A comprehensive specification of the models and their parameters is given in **Supplement S5**.

### General model assumptions

First, each feature of a stimulus in the world is represented by a sensory measurement *m*′ in the observer’s brain. Based on previous work [27, 63, 64], we assumed the measurements might be biased. Thus, the average measurement equals a remapped feature value *s*′, which is a linear function of the physical feature value *s*.

Second, the measurements are corrupted by Gaussian noise *m*′ ∼ 𝒩(*s*′, *σ*′^2^). The standard deviation of repeated measurements across trials reflects the actual sensory uncertainty *σ*′associated with the stimulus (**Fig. 1B**).

Third, the observer has prior knowledge (or prior assumptions) about the probability distribution of the stimulus features. For example, the observer might assume that stimuli most likely are located straight ahead, 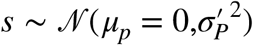 [49].

Fourth, in every trial, the observer derives the posterior probability of a stimulus feature by combining the prior and the likelihood. Because the sensory noise is Gaussian, the likelihood is a Gaussian function centered on the measurement. The standard deviation of the likelihood corresponds to the estimated uncertainty, 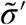, which might differ from the actual sensory uncertainty, 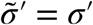 or 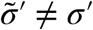 (**Fig. 1C**). The two estimated uncertainties we consider that might differ from actual sensory uncertainty are for auditory spatial-location measurements 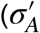 vs 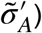 and audiovisual SOA measurements 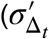 vs 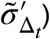.

Fifth, the observer uses the mode of the posterior as the (conditional) estimate of the stimulus feature.

Sixth, the task-relevant estimates of the stimulus feature(s) are used to generate a perceptual decision. Depending on the task, perceptual decisions might be based directly on one or two feature estimates or might involve further inferences such as those over the causal scenario underlying auditory and visual measurements.

Finally, the response indicating the perceptual decision is subject to additional noise sources such as motor noise and attentional lapses.

### Model variants

We tested four models: Either none, one, or both of the uncertainty estimates, 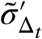 or 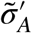, were independent of the actual uncertainty 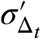 or 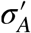. We fit each model jointly to the data from all four experiments. For example, if in a model perceptual inferences were based on an accurate estimate of auditory spatial uncertainty 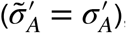, this held for the modeling of all inferences that used an estimate of auditory spatial uncertainty. Specifically, Expt. 1 constrained 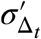, Expts. 2 and 3 constrained 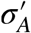 and 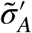, and Expt. 4 constrained 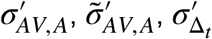 and 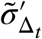. Based on previous results [55, 65], we assumed that auditory spatial uncertainty might differ in unimodal (Expts. 2 and 3) compared to bimodal contexts (Expt. 4, so that 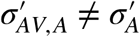.

### Model fitting

The models of the five tasks were fit jointly; the log-likelihood of the full parameter set (see **Supplement Tables S4-S6**) was calculated by combining the log-likelihoods of each task’s model and its parameters. The best-fitting parameters, for which the joint likelihood was maximal, were derived using the BADS toolbox [66]. This joint fit was repeated for each of the four model variants. To compare model performance across model variants, we computed the Akaike information criterion [AIC; 67] separately for each participant. The winning model corresponds to the model with the minimal AIC value. We compared the best-fitting parameter values of the actual and employed audiovisual temporal and auditory spatial uncertainty using pairwise *t*-tests.

## Results

To identify whether participants employed correct estimates of their temporal and spatial sensory uncertainty for perceptual inference, we contrasted a model that assumed participants’ perceptual decisions relied on accurate estimates of their audiovisual temporal and auditory spatial uncertainty with models that allowed for inaccurate estimates of uncertainty. We further distinguished models that assumed inaccurate temporal uncertainty, inaccurate spatial uncertainty, or both.

Each model was jointly fit to a participant’s responses in five tasks tested in four separate experiments: (1) audiovisual temporal-order judgment, (2) audiovisual spatial-discrimination, (3) auditory and visual spatial-localization, (4) auditory spatial-localization for auditory stimuli accompanied by a temporally and/or spatially discrepant visual stimulus and (5) a causal-inference task in which participants determined the number of sources of these audiovisual stimulus pairs (see **Fig. 3** and Methods).

The behavioral results in all experiments showed clear signatures of task-specific perceptual inference, known to rely on estimates of the uncertainty associated with a sensory measurement (see **Fig. 4** for the data of a representative participant and **Supplement S3** for all participants). Responses in the audiovisual spatial-discrimination task (Expt. 2) showed clear cross-modal biases (**Fig. 4B**) that were also reflected in the localization of auditory (**Fig. 4C**) but not visual (**Supplement S3.3.1 - 3.3.2**) stimuli in the unimodal localization task (Expt. 3). In the bimodal context of Experiment 4, auditory localization responses were shifted towards the accompanying visual stimulus, indicative of multisensory integration (**Fig. 4D**). Moreover, these perceptual shifts as well as the proportion of trials in which participants perceived the auditory and the visual stimulus as sharing a common cause decreased with increasing spatial and temporal discrepancy, demonstrating that participants performed causal inference (**Figs. 5A,B** top panels; see **Supplement S3.4** for statistical analysis).

**Figure 4.**
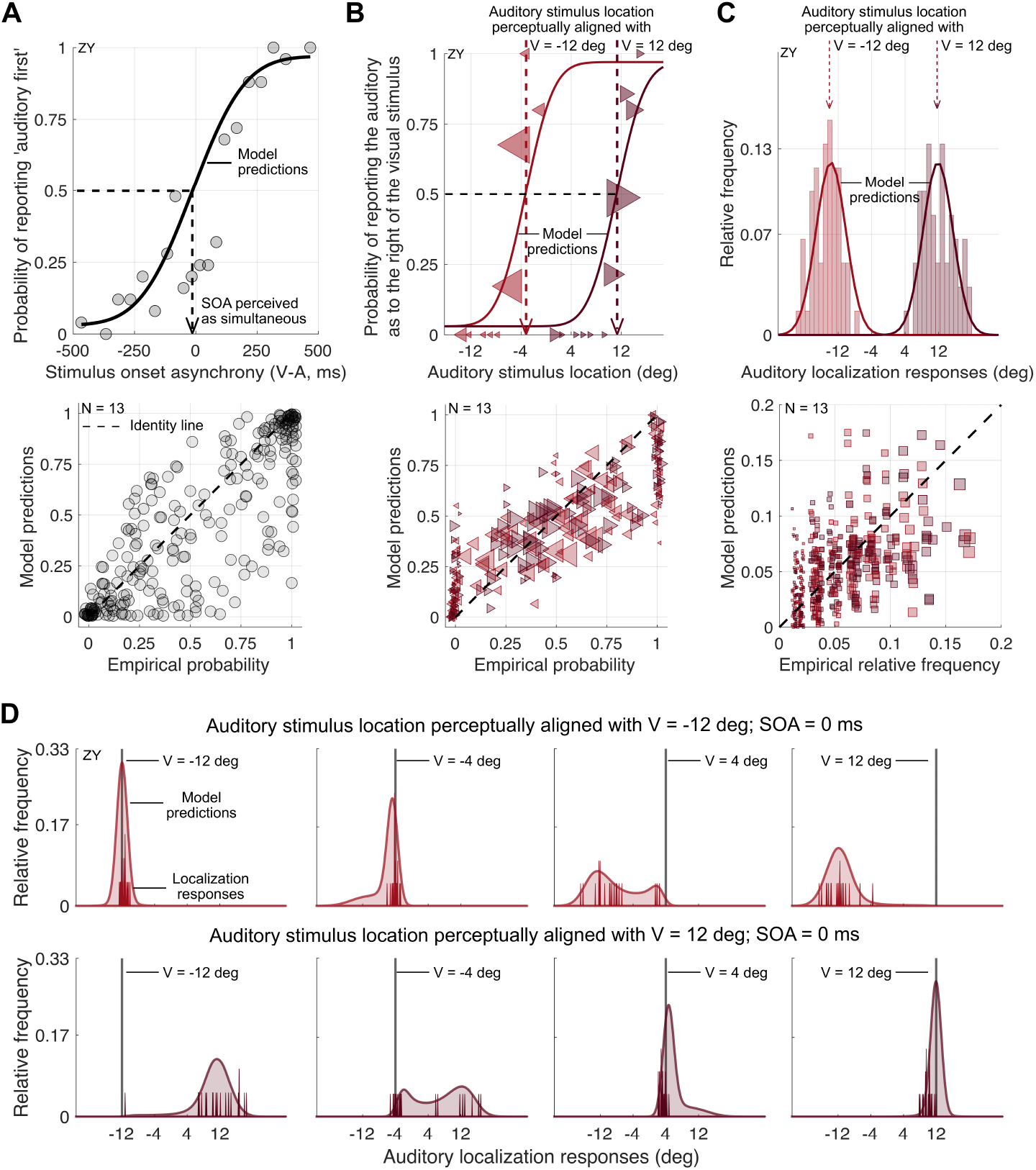
Behavioral results and joint model predictions for audiovisual temporal and spatial discrimination tasks as well as auditory localization in unimodal and bimodal contexts. (A) Expt. 1 (audiovisual temporal-order judgment). Top panel: observed (gray dots) and predicted (solid line) proportions of auditory-came-first reports as a function of stimulus onset asynchrony (SOA) of the auditory and visual stimulus (negative values: visual first) for an example participant. Bottom panel: probability of reporting auditory-first, empirical data versus model predictions for all participants and SOAs. (B) Expt. 2 (audiovisual spatial discrimination). Top panel: observed proportion of trials in which the auditory stimulus was reported as located to the right of the visual one as a function of the binned auditory stimulus location (markers, area proportional to the number of trials) for the same participant. Observed proportions and model predictions (lines) are shown separately for trials in which the visual stimulus was presented at -12 deg (leftward-pointing light red triangles) and trials in which the visual stimulus was presented at 12 deg (rightward-pointing dark red triangles). Dashed arrows: auditory stimulus locations perceptually aligned with visual stimuli at -12 and 12 deg. These auditory stimulus locations were used in all other experiments. Bottom panel: probability of reporting the auditory stimulus as to the right of visual one, (binned) empirical data versus model predictions for all participants. Marker size is proportional to the number of trials. (C) Expt. 3 (auditory localization in unimodal contexts). Top panel: histogram of localization responses and model predictions (lines) for the same participant. The auditory stimulus was either presented at the location corresponding to a visual stimulus at -12 deg (light red) or to a visual stimulus at 12 deg (dark red). Auditory and visual stimuli were interleaved (see **Supplement S3** for data from the visual-localization task). Bottom panel: binned observed versus predicted relative frequencies of localization responses for all participants. (D) Expt. 4A (auditory localization in audiovisual contexts). Observed (solid vertical red lines) and predicted (shaded area) frequencies of localization responses for auditory stimuli presented as part of an audiovisual stimulus pair with a SOA of 0 ms for the same participant. The auditory stimulus was either presented at a location perceptually aligned with a visual stimulus at -12 deg (top row) or 12 deg (bottom row). The visual stimulus was presented at one of four locations (±4, ±12 deg; columns). See **Supplement S3** for data and model predictions for all audiovisual SOAs.

**Figure 5.**
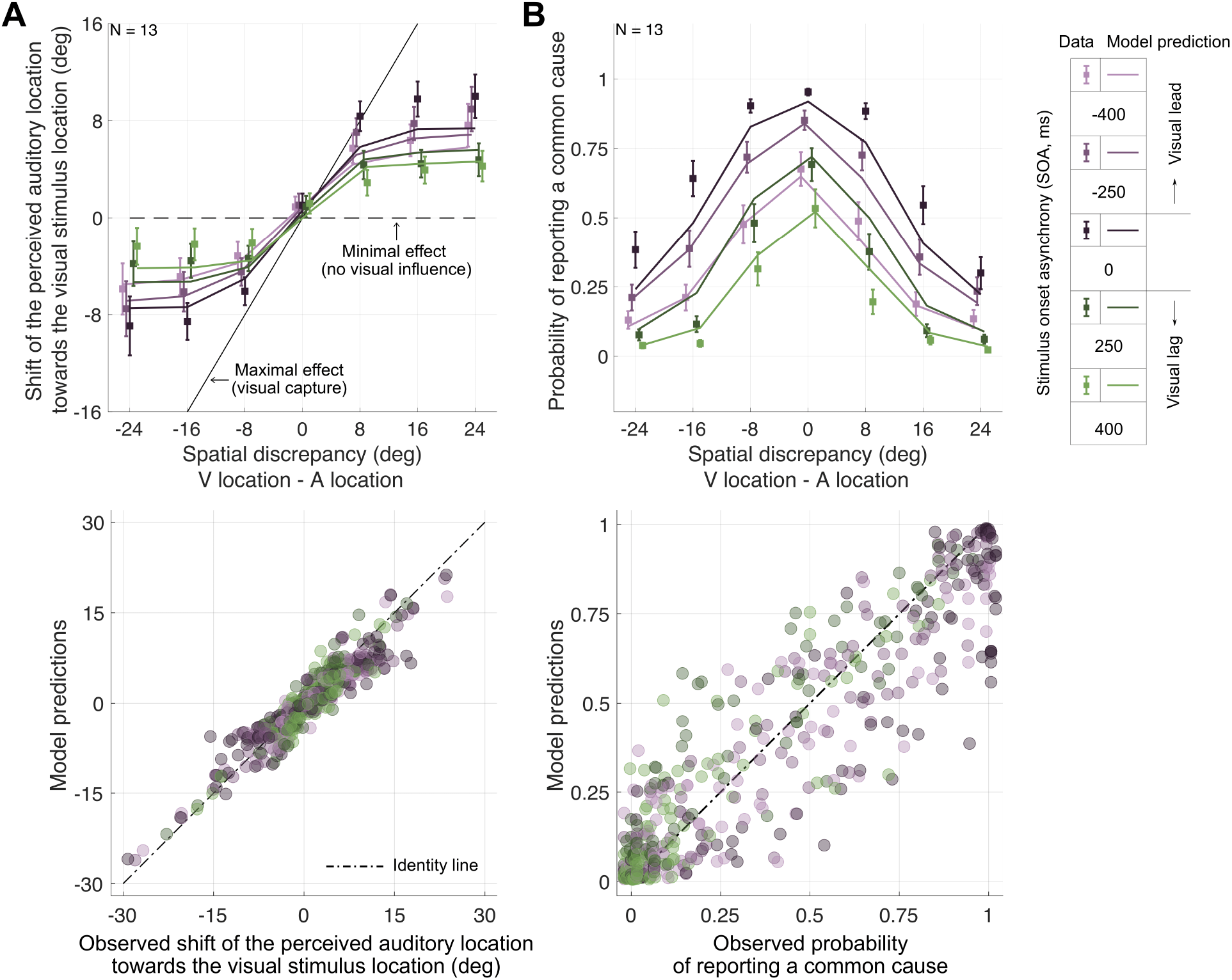
Behavioral results and model predictions for auditory localization in bimodal contexts and audiovisual causal-inference judgments. (A) Top panel: Group average shifts of perceived auditory location towards the visual stimulus location for each audiovisual temporal discrepancy as a function of audiovisual spatial discrepancy. Data jittered slightly horizontally for legibility. Bottom panel: observed versus predicted shifts in auditory localization responses towards the visual location. (B) Probability of reporting a common cause as a function of spatial discrepancy for each temporal discrepancy (colors). Markers: group averages, error bars: ±1 SEM. Bottom panel: observed versus predicted proportion of common-cause judgments.

We assumed that parameters that influenced perceptual decisions did so consistently across all five tasks. Thus, we modeled data from these five tasks jointly, assuming consistent parameter values across tasks. Jointly fitting a set of observer models to all of the data allowed us to simultaneously constrain parameters of interest, those reflecting the participant’s actual and employed temporal and spatial auditory uncertainty, as well as additional parameters that might trade off against parameters of interest, including biases in auditory spatial perception, biases in audiovisual temporal perception, and prior assumptions regarding the prevalence of some spatial locations and temporal discrepancies. The validity of this approach is demonstrated in the general agreement between data and model predictions across all tasks (**Figs. 4A-C**, bottom panels; **Figs. 5A,B**, bottom panels).

The behavioral results were best captured by a model variant that allowed perceptual inferences to be based on inaccurate estimates of both temporal and spatial uncertainty (11 out of 13 participants, see **Supplement S6.2** for measures of model fit for each model variant and participant and for model comparisons). The behavior of the two remaining participants was best captured by model variants that assumed that only inaccurate temporal or inaccurate spatial uncertainty estimates were employed. Parameter estimates of the winning model reveal how gravely and consistently participants underestimated their sensory uncertainty, that is, by how much the uncertainty estimates participants used for perceptual decisions differed from their actual sensory uncertainty. The temporal uncertainty estimates participants’ employed for inference (median = 74.93 ± 14.02 ms) were about half of actual temporal uncertainty (median = 157.68 ± 14.02 ms; *t*(12) = 4.3, *p* = 0.001; **Fig. 6A**). The same mismatch was observed for auditory spatial uncertainty in unimodal and bimodal contexts: the employed uncertainty was less than half of actual uncertainty (**Fig. 6B,C**; unimodal context: median = 1.46 ± 0.41 deg vs. 5.45 ± 0.92 deg; *t*(12) = 4.79, *p* < 0.001; bimodal context: median = 7.18 ± 4.91 deg vs. 15.58 ± 1.93 deg; *t*(11) = 2.79, *p* = 0.018). These results provide clear evidence that humans’ inferences rely on overconfident estimates of audiovisual temporal and auditory spatial uncertainty. Finally, to get a feel for how the model leads to these predictions, see the results of model simulations as parameters are varied in **Supplement S4**.

**Figure 6.**
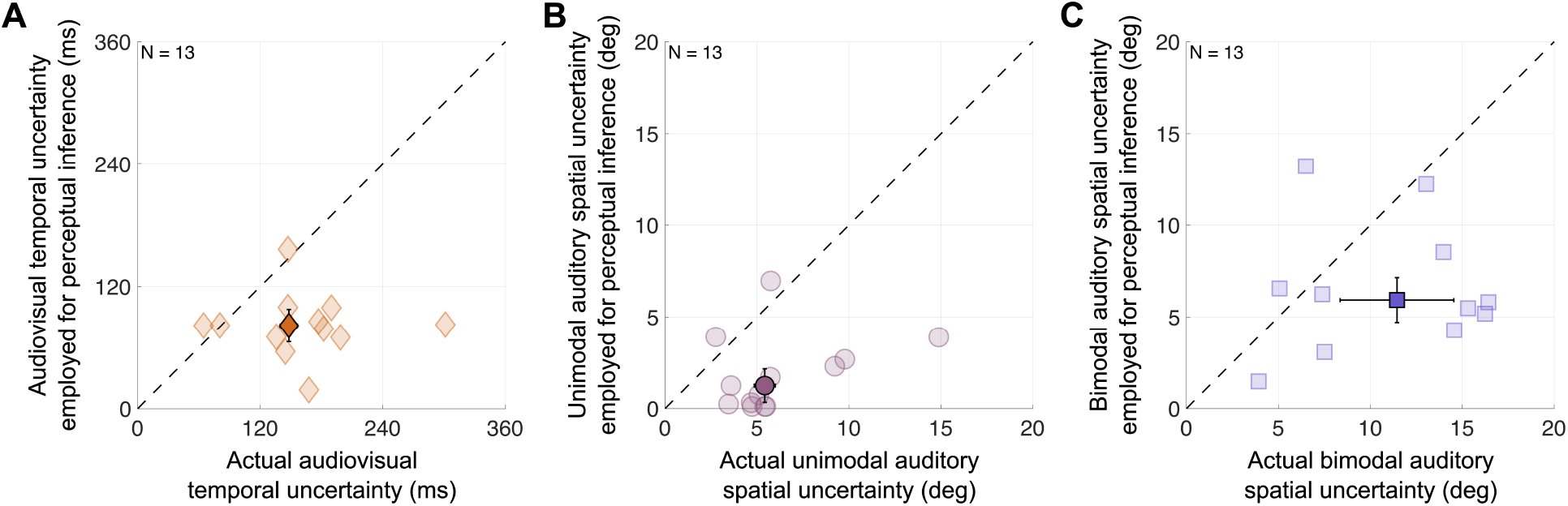
Biases in estimates of auditory spatial and audiovisual temporal uncertainty. Best-fitting uncertainty parameters of the winning model allowing for perceptual inferences based on inaccurate estimates of temporal and spatial uncertainty. Participants’ actual sensory uncertainties (the standard deviations of the measurement distributions) are plotted against the uncertainty values they used for perceptual inference (the standard deviations of the likelihood functions). (A) Audiovisual temporal uncertainty. (B) Auditory spatial uncertainty given an auditory stimulus presented alone or (C) accompanied by a visual stimulus. Darker markers: group medians. Error bars: SEM. Dashed lines: identity. One outlier is not displayed in (C) to prevent excessive scaling of the axes (see **Supplement S6**).

## Discussion

In this study, we tested the ubiquitous assumption that perceptual inference relies on accurate estimates of sensory uncertainty. Participants completed a series of experiments encompassing audiovisual temporal-order judgments, audiovisual spatial discrimination, auditory spatial localization in unimodal and bimodal contexts, as well as causal-inference judgments. Perceptual decisions in these experiments take the observer’s auditory and visual spatial uncertainty as well as their audiovisual temporal uncertainty into account. To evaluate whether participants’ perceptual inferences relied on accurate estimates of these sensory uncertainties, we contrasted models that differed in the agreement between actual and employed temporal and spatial uncertainty. Data from all tasks were fit jointly to constrain estimates of all components of perceptual inference. Model comparisons and evaluation of the best-fitting parameters revealed that the majority of participants systematically underestimated audiovisual temporal and auditory spatial uncertainty. This pattern held across unimodal and bimodal contexts, indicating that the revealed mismatch between actual and employed sensory uncertainty is a general phenomenon.

The results revealed that observers used overconfident estimates of auditory spatial and audiovisual temporal uncertainty for various perceptual inferences. In contrast, visual spatial uncertainty — in accordance with previous studies [25] — was assumed to be accurately represented, suggesting that the quality of uncertainty estimates varies across modalities. This discrepancy might be rooted in fundamental differences between the encoding of spatial location in primary visual and auditory cortices. Bayesian theories of neural coding propose that sensory uncertainty is automatically encoded in the population response of neurons that are tuned to the feature of interest [12]. In the visual system, various features such as location, orientation, and motion direction are encoded by tuned populations [11, 68, 69], rendering such a mechanism possible. In contrast, spatial location in the auditory cortex is coded by opponent neural populations that are broadly tuned for far-left or far-right locations [70, 71]. Thus, models that describe the neural representation of uncertainty for visual features cannot be easily extended to auditory spatial uncertainty.

It seems unlikely that a probabilistic population code is used for temporal discrepancy as there is no specific sensory system specialized for time perception. Neuroimaging research suggests that temporal-order information is processed in a distributed network of brain regions across prefrontal, fronto-parietal, parietal, and occipito-temporal cortices [72-74]. Given that multiple brain areas are involved when encoding cross-modal temporal discrepancy, the representation of sensory uncertainty might be similarly distributed and difficult to decode. Estimation of temporal uncertainty might require combining information across these regions, which could lead to a mismatch between actual and employed temporal uncertainty.

We observed consistent underestimation of sensory uncertainty across participants, tasks and domains, which puts the costs and benefits of such a misestimation into focus. Simulation results suggest that underestimating sensory uncertainty improves accuracy at the expense of precision. However, the tradeoff between accuracy and precision appears to be beneficial only when the internally remapped stimulus location closely matches the physical stimulus location. The presence of biases in auditory spatial perception undermines the possibility of achieving accuracy through overconfident sensory uncertainty. Nonetheless, perceptual biases are inherent and likely to remain undetected by individuals [63].

The brain constantly makes decisions based on noisy sensory information. The extent to which the brain employs an accurate estimate of sensory uncertainty has profound implications for perception, action, and cognition. Sensory uncertainty is a critical variable in a myriad of models describing human behavior across a wide range of domains, from perceptual inference [75, 76] to cognitive and metacognitive judgment [77-79] to sensorimotor control [80]. Central to these models is the assumption that inference relies on accurate estimates of sensory uncertainty. Our findings suggest a critical revision: models need to include the possibility that sensory uncertainty is not accurately represented. The spread of the likelihood function that underlies an inference might not match that of the distribution of noisy measurements across trials. Incorporating this mismatch introduces greater model complexity, increasing the risk of overfitting by adding more degrees of freedom. To mitigate this risk, researchers should emphasize experimental designs ensuring that all parameters are properly constrained.

In conclusion, this study revealed that perceptual inference in different contexts relies on overconfident estimates of auditory spatial and audiovisual temporal sensory uncertainty.

