## Supplemental materials for "Humans use overconfident estimates of auditory spatial and temporal uncertainty for perceptual inference"

**S1: Detailed experimental procedures** **3**
S1.1: Experiment 1 - Temporal-order-judgment task 3
S1.2: Experiment 2 - Audiovisual spatial-discrimination task 3
S1.3: Experiment 3 - Unimodal spatial-localization task 6
S1.4: Experiment 4 - Auditory spatial-localization task in audiovisual contexts and explicit causal-inference judgments 6
**S2: Participant-specific auditory stimulus locations** **8**
**S3: Participant-level results and model predictions** **10**
S3.1: Experiment 1 - Temporal-order-judgment task 10
S3.2: Experiment 2 - Audiovisual spatial-discrimination task 11
S3.3: Experiment 3 12
S3.3.1: Localization practice 12
S3.3.2: Unimodal visual-localization task 13
S3.3.3: Unimodal auditory-localization task 14
S3.4: Experiment 4 15
S3.4.1: Auditory-localization task in the bimodal context 15
S3.4.2: Causal-inference judgments 17
**S4: Simulation results** **19**
**S5: Modeling** **22**
S5.1: Formalization of the audiovisual temporal-order-judgment task (Experiment 1) 22
S5.2: Formalization of the audiovisual spatial-discrimination task (Experiment 2) 23
S5.3: Formalization of the unimodal auditory and visual spatial-localization tasks (Experiment 3) 24
S5.4: Formalization of the auditory spatial-localization task in a bimodal context (Experiment 4A) 25
S5.5: Formalization of the causal-inference task (Experiment 4B) 28
S5.6: Model variants 29
S5.7: Model likelihoods 31
S5.7.1: Model likelihoods for the audiovisual temporal-order-judgment task (Experiment 1) 31
S5.7.2: Model likelihoods for the audiovisual spatial-discrimination task (Experiment 2) 32
S5.7.3: Model likelihoods for the unimodal auditory and visual spatial-localization task 33
S5.7.4: Model likelihoods for the bimodal spatial-localization and causal-inference tasks 34
S5.8: Model fitting 36
S5.8.1: Parameter estimation 36
S5.8.2: Model comparison 36
**S6: Model-estimated uncertainties and model comparison** **37**
S6.1: Model-estimated actual and employed uncertainties for perceptual inference 37
S6.2: Model comparison 38
**References** **40**

### S1: Detailed experimental procedures

Participants completed four experiments across three separate sessions, spanning three days. These included: (1) an audiovisual temporal-order-judgment task (2) an audiovisual spatial-discrimination task, (3) a unimodal spatial-localization task, and (4) an auditory spatial-localization task in audiovisual contexts followed by causal-inference judgments. On the first day, participants completed Experiments 2 and 3. The second day was dedicated to completing two-thirds of Experiment 4, while the final day covered the remaining third of Experiment 4 and Experiment 1.

#### S1.1: Experiment 1 - Temporal-order-judgment task

In this two-alternative, forced-choice task, participants judged the temporal order of audiovisual stimulus pairs with varying SOA. Each trial started with a fixation cross presented straight ahead for 500 ms, followed by 500 ms of blank screen. Then, a temporally offset stimulus pair, an auditory and a visual stimulus each 100 ms-long, was presented straight ahead, followed by 700 ms of blank screen. Next, a response probe appeared on the screen, and participants reported by button press which stimulus they perceived first, the auditory or the visual one (**Fig. S1A**). There was no time constraint for the response. Response feedback was not provided. The inter-trial interval was 500 ms.

The SOA between the auditory and the visual stimuli ranged from -366.67 to 366.67 ms (positive: auditory-leading; negative: visual-leading). The interval steps varied: from -366.67 to -83.33 ms, the step size was 50 ms; from -83.33 to 83.33 ms, the step size was 33.33 ms; and from 83.33 to 366.67 ms, the step size increased back to 50 ms. We additionally included two very long SOAs, -466.67 and 466.67 ms to better estimate the lapse rate, resulting in a total of 20 SOAs. Each SOA was tested 25 times, resulting in a total of 500 trials administered in pseudorandom order. These trials were split into five blocks; participants were encouraged to rest in between blocks for as long as they desired. Usually participants took an hour to complete this task.

#### S1.2: Experiment 2 - Audiovisual spatial-discrimination task

In this two-alternative, forced-choice task, participants judged the relative location of an auditory compared to a visual stimulus. Specifically, in each trial, participants were presented with one of two visual standard stimuli in one interval and an auditory test stimulus in the other interval, in pseudo-randomized order. Participants then judged the location of the auditory stimulus relative to that of the visual standard stimulus. Each trial began with a fixation cross, presented at the center of the screen for 500 ms, followed by a blank screen lasting 500 ms. Then, the first 100 ms-long stimulus was presented, the second stimulus followed with an SOA of 1600 ms. After another 500 ms blank screen, a response probe appeared, and participants responded by button press whether they perceived the auditory stimulus as to the right or to the left of the visual one. Feedback was not provided (**Fig. S1B**).

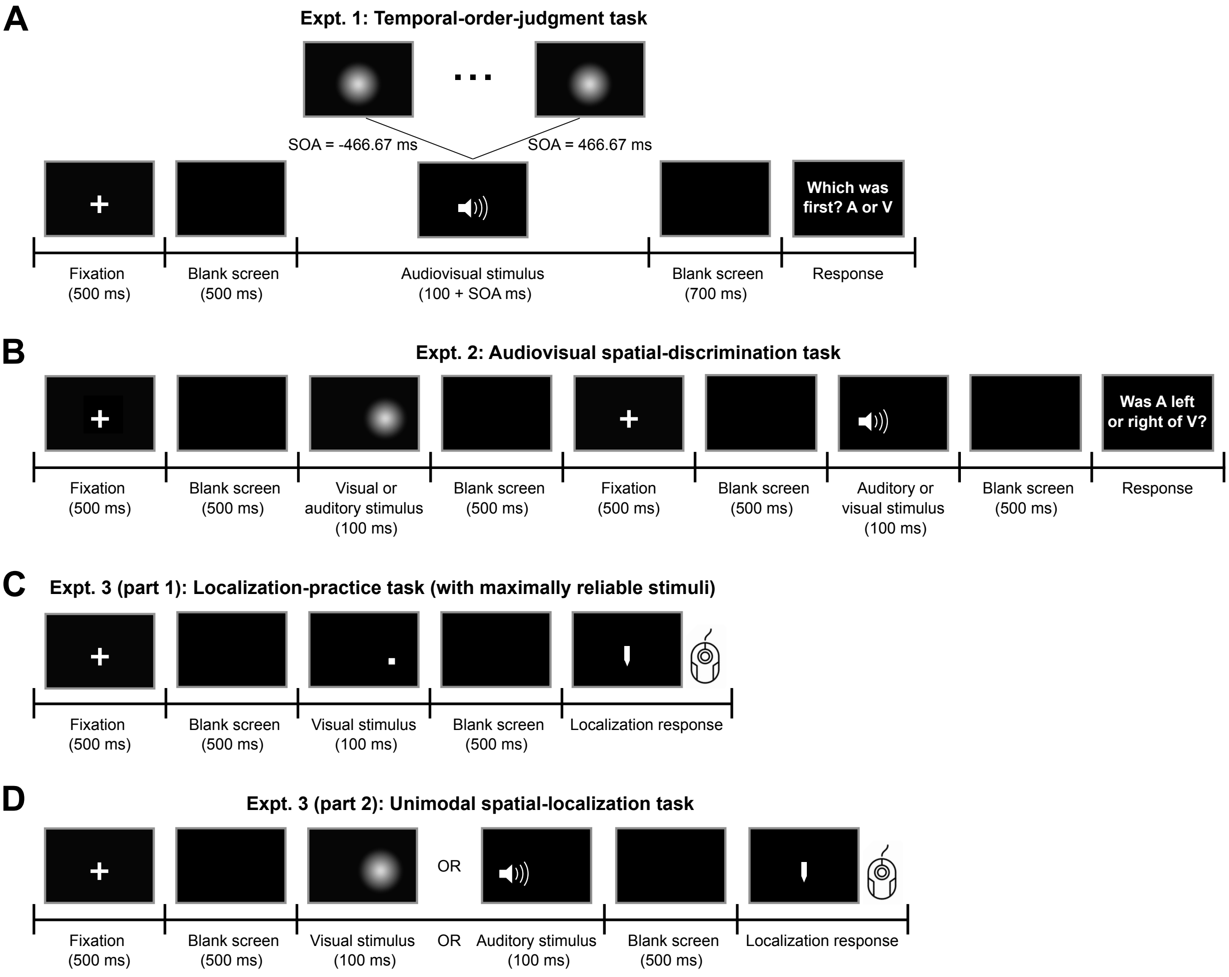


**Figure S1. Trial events sequence in Expts. 1-3.** (A) Temporal-order judgment task – Experiment 1. Participants were presented with an audiovisual stimulus pair with varying stimulus onset asynchrony (SOA, negative values indicate visual-leading stimulus pairs). Each element (auditory or visual) of the audiovisual stimulus pair lasted for 100 ms. Participants reported which stimulus they perceived first by button press. (B) Audiovisual spatial-discrimination task – Experiment 2. Participants were presented with a visual standard in one (randomly selected) interval and an auditory test stimulus in the other interval. They reported whether they had perceived the auditory stimulus as to the left or right of the visual stimulus. (C) Task timing of the unimodal visual spatial-localization task (the localization-practice task with maximally reliable stimuli). Participants were presented with a white square and used a visual cursor to indicate the stimulus' horizontal location. (D) Unimodal spatial-localization task – Experiment 3, part 2. Participants localized either a visual or an auditory stimulus by moving a visual cursor to the horizontal location of the stimulus.

After the response, the loudspeaker moved to its new location. The movements of the speaker were audible, and we were concerned that participants might infer the position of the speaker from that sound. To mask any auditory cues about the new location of the speaker, a masking sound was played during this period. The masking sound (55 dB) consisted of a recording of the sound generated by a randomly chosen speaker movement plus white noise. To further foil the use of auditory cues from the moving sledge, every time the speaker moved to a new target location, it first moved to a stopover location. The stopover location was randomly chosen under the constraint that the total distance the speaker moved and the amount of time

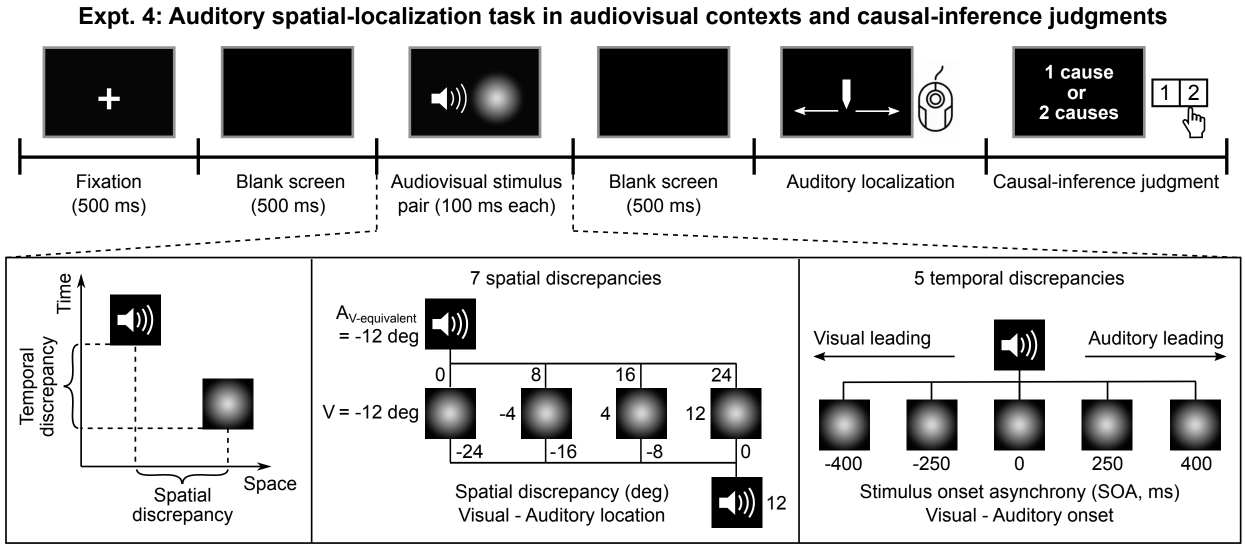


**Figure S2. Trial events sequence and stimuli in Experiment 4.** Participants were presented with an audiovisual stimulus pair with varying spatial ($\pm 24$, $\pm 16$, $\pm 8$ and $0$ deg computed using visual physical locations and auditory locations with the perceptually equivalent visual scale) and temporal discrepancies ($\pm 400$, $\pm 250$ and $0$ ms). After the stimulus presentation, they sequentially completed two tasks: (1) they localized the auditory element using a visual cursor, and (2) they reported by button press whether they perceived the two stimuli as originating from a common source or two separate sources. The audiovisual stimulus pairs varied with respect to the temporal and spatial discrepancy between the two elements. Auditory stimulus locations were chosen in perceptual space based on Experiment 2 (see **S2** for more details).

the movement took were approximately equal across trials. The combinations of these two methods proved effective in eliminating location cues (1).

The visual stimulus was presented at either -12 or 12 deg relative to the center of the screen. The location of the auditory test stimulus was controlled by four interleaved staircases, two for each visual stimulus location. Of the two staircases, one started the auditory test stimulus to the left of the visual standard stimulus, and the other to the right side of the visual standard stimulus (for $V$ = -12 deg, $A_{\text{left}}$ = -13.25 deg and $A_{\text{right}}$ = 1.25 deg; for $V$ = 12 deg, $A_{\text{left}}$ = -1.25 deg and $A_{\text{right}}$ = 13.25 deg). Staircases starting from the right side followed the one-down-two-up rule, converging to a probability of 29% of choosing the auditory stimulus as farther to the right than the corresponding visual standard stimulus. Staircases starting from the left side followed the two-down-one-up rule, converging to a probability of 71% of choosing the auditory stimulus as farther to the right than the corresponding visual standard stimulus. The initial step size was 1.9 deg, which was decreased to 1.0 deg after the first staircase reversal and to 0.5 deg after the third reversal. Each staircase comprised 36 trials. An easy trial in which the auditory test stimulus was presented at either one of the two start locations (e.g., -13.25 or 1.25 deg for a visual standard of -12 deg) was inserted once every 9 trials to improve lapse-rate estimation, resulting in a total of 160 trials. The experiment was divided into four blocks. Usually, participants took about an hour and a half to complete this task.

#### S1.3: Experiment 3 - Unimodal spatial-localization task

At the beginning of this experiment, participants practiced using a rollerball mouse to adjust the visual cursor while localizing a maximally reliable visual stimulus. Each trial started with a central fixation cross, which was presented for 500 ms, followed by a 500 ms blank screen. Then, the visual training stimulus, a small high-resolution white square, was displayed on the screen for 100 ms, followed by a blank screen, presented for 500 ms. Next, a response cursor appeared at a random horizontal location on the screen. Participants moved the response cursor to the location of the stimulus by moving the rollerball, and clicked one of the mouse keys to register the response (**Fig. S1C**). The training stimulus was presented in one of eight locations, evenly spaced from -17.5 to 17.5 deg in steps of 5 deg. Each stimulus location was visited 30 times in random order, resulting in a total of 240 trials. The inter-trial interval was 500 ms. This experiment usually took half an hour to complete.

After the practice, participants performed a unimodal spatial-localization task on the experimental stimuli used in all regular tasks. Each trial started with a fixation cross presented straight ahead for 500 ms, followed by 500 ms of blank screen. Then, either an auditory or a visual stimulus was presented for 100 ms, followed by 500 ms of blank screen. Next, the response cursor appeared in a random location, and participants adjusted the horizontal location of the cursor to match that of the stimulus (**Fig. S1D**). There was no time limit for the response. Visual feedback of the cursor location was provided during adjustment, but localization-error feedback was not provided. After the response, the loudspeaker moved to its new location. To eliminate location cues, we again moved the speaker to a stopover position and played the masking sound during the movements of the speaker (see **Supplement S1.2**). The inter-trial interval was kept constant across visual and auditory stimuli and due to the necessary speaker movement was approximately 3000 ms. The visual stimulus was presented at either one of four locations (±4, ±12 deg relative to straight ahead); each visual stimulus location was tested 20 times. The auditory stimulus was presented at one of two participant-specific locations, identified as perceptually aligned with visual stimuli at ±12 deg in Expt. 2; see section **S2** for further information. Each auditory stimulus location was tested 40 times. The unimodal visual and auditory stimuli were interleaved, resulting in a total of 160 trials administered in pseudorandom order. These trials were split into four blocks. Usually participants took an hour to complete this task.

#### S1.4: Experiment 4 - Auditory spatial-localization task in audiovisual contexts and explicit causal-inference judgments

In this task, participants were presented with audiovisual stimulus pairs with various spatial and temporal discrepancies (**Fig. S2**). Each trial started with the presentation of a fixation cross at the center of the screen for 500 ms, followed by 500 ms of blank screen. An audiovisual stimulus pair was presented, followed by 500 ms of blank screen. Next, the response cursor appeared, and participants used the rollerball mouse to adjust the horizontal location of the cursor to localize the auditory component. After the localization response, a response prompt for the explicit causal-inference task (also known as a unity judgment) was displayed. Participants reported whether they perceived the auditory and the visual stimulus as coming from a common source or from two separate sources using the mouse keys (left click = common source, right click = two sources). After the response, the loudspeaker moved to its new location. To eliminate location cues, we again moved the speaker to a stopover position and played the masking sound during the movements of the speaker (see **Supplement S1.2**).

The visual stimulus was presented at ±4 or ±12 deg; the auditory stimulus was presented at one of the same two locations as in Expt. 3, those at which the auditory stimuli were perceptually aligned with visual stimuli at ±12 deg. The stimuli were presented with an SOA of 0, ±250, or ±400 ms (negative values indicate visual first). There were 40 different audiovisual stimulus pairs (4 visual stimulus locations x 2 auditory stimulus locations x 5 SOAs). Each of these pairs was tested 20 times, resulting in a total of 800 trials, presented in a pseudo-randomized order. These trials were split into two sessions. Participants completed the first session (560 trials) typically in two hours on one day, and completed the second session (240 trials) in approximately an hour on another day.

#### S2: Participant-specific auditory stimulus locations

One additional goal of Expt. 2 was to identify auditory locations that were perceived as co-located with two pre-selected visual locations for each participant. Presenting auditory stimuli at these locations allowed us a more precise investigation of perception in audiovisual contexts (Expt. 4). The following analysis was performed right after the participant completed the task, as its outcome determined locations of auditory stimuli that were used in subsequent tasks. The joint model fits (**S3**) were conducted independent of the analysis outlined below.

**Analysis**. Responses were coded as a binary variable indicating whether the auditory test stimulus had been perceived as located to the right of the visual standard stimulus. We fitted two cumulative Gaussian distributions to these data as a function of auditory stimulus location, one curve for each visual standard stimulus location, with a common lapse rate, constrained to be less than 6% (2). The point of subjective equality (PSE) was defined as the auditory stimulus location corresponding to a probability of 0.5 of reporting the auditory stimulus to the right of the visual stimulus on the psychometric function (**Fig. S3A**; see **Fig. S4** for all participants' data). We computed a PSE from each fitted psychometric curve, i.e., for each visual stimulus location we determined the location at which an auditory stimulus was perceptually co-located with the visual stimulus. To derive error bars for the PSEs, separately for each visual standard stimulus, we randomly resampled the raw data with replacement 1,000 times, fitted cumulative Gaussian distributions to each resampled dataset, computed the PSEs, and took the 2.5th and 97.5th percentiles of the 1,000 PSEs as the bootstrapped confidence interval.

**Results**. The group-averages of the auditory stimulus locations that perceptually matched the two pre-selected visual stimulus locations (-12 and 12 deg, respectively) were -6.71 deg (SEM = ±0.52 deg) and 7.59 deg (SEM = ±0.85 deg) (**Fig. S3B**), indicating that the auditory stimuli were perceived as shifted toward the periphery relative to the visual stimuli, which is in line with previous findings (3-5). These participant-specific auditory locations were used in Experiments 3 and 4, to ensure minimal spatial discrepancy in perceptual space when auditory stimuli were paired with visual stimuli presented at ±12 deg (**Fig. S3C**).

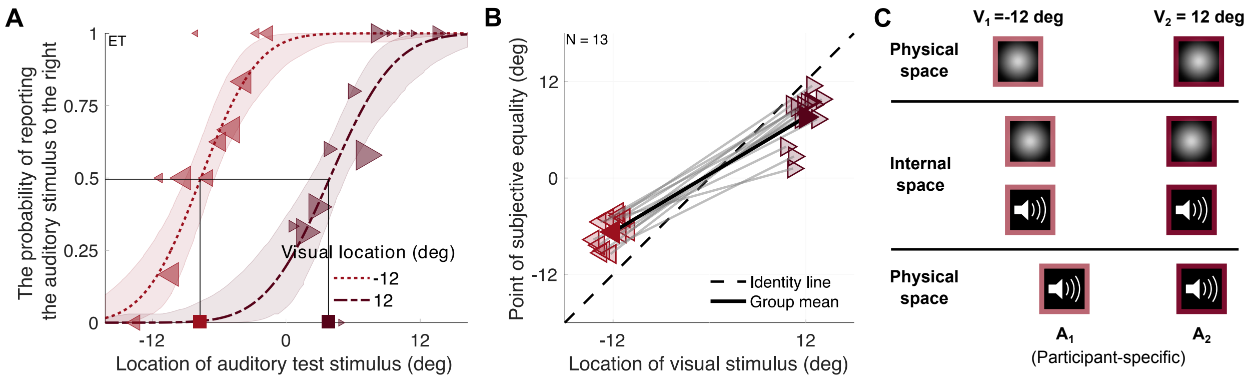


**Figure S3. Experimental procedure and behavioral results for preparatory Expt. 1.** (A) Behavioral results of representative participant ET. Two cumulative Gaussian functions (solid lines: best-fitting curves; shaded areas: 95% bootstrapped confidence intervals), one for each visual standard stimulus, are fitted to psychometric data (dots, bin size  1 deg; marker area proportional to the number of trials in each bin). The point of subject equality (PSE) was computed for each visual standard stimulus (squares). (B) PSE as a function of visual standard stimulus location. Dashed black line: identity line; black solid line: group average; gray solid lines: individual data. (C) Stimulus locations in physical and perceptual space. Top panel: visual standard stimuli in physical space; middle panel: the two pairs of a visual and an auditory stimulus that were aligned in perceptual space; bottom panel: the two participant-specific auditory stimuli in physical space.

### S3: Participant-level results and model predictions

#### S3.1: Experiment 1 - Temporal-order-judgment task

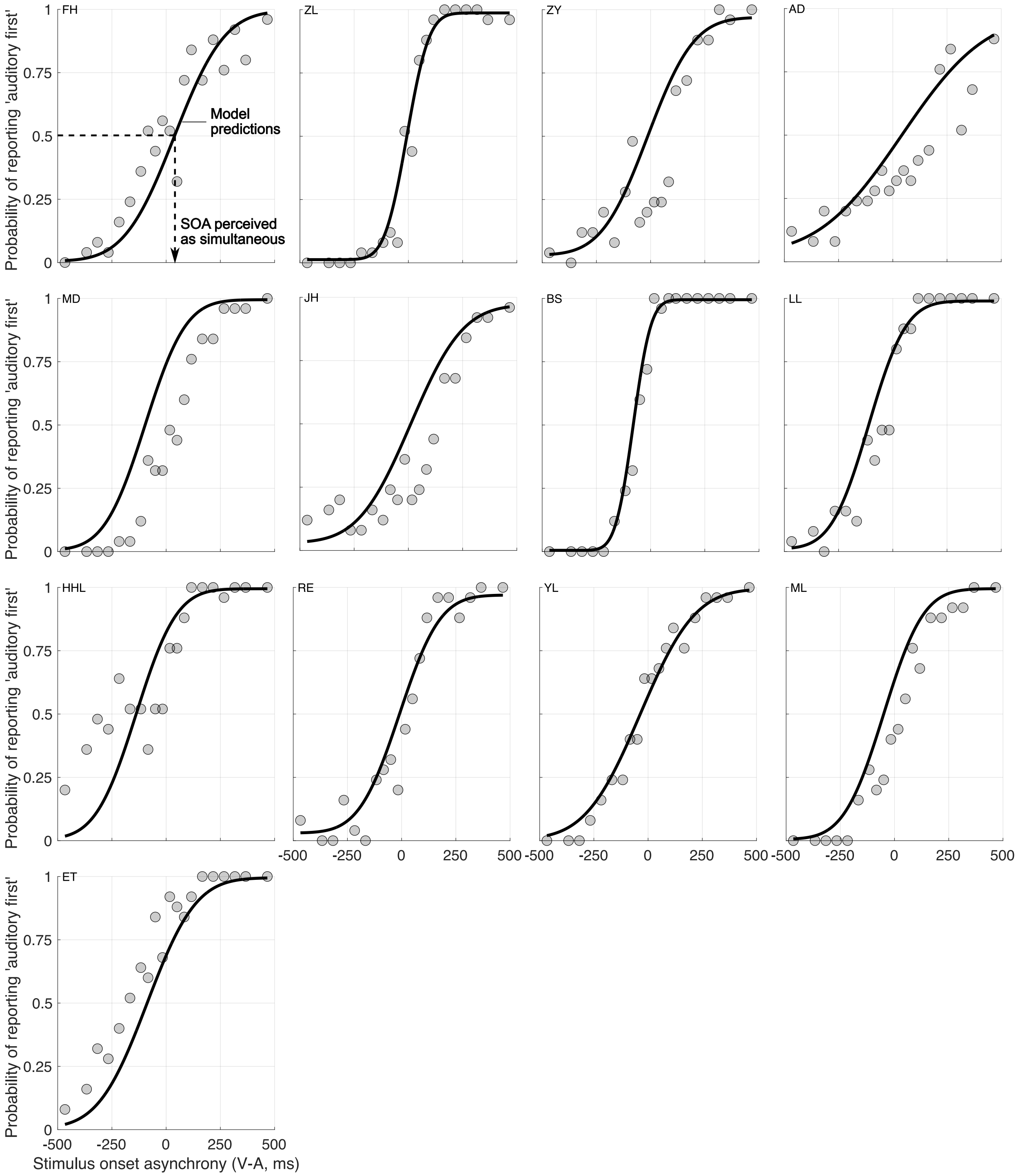


**Figure S4. Results of the temporal-order-judgment task for all participants.** Participants indicated the temporal order of audiovisual stimulus pairs. Observed (gray dots) proportions of auditory-came-first reports are plotted as a function of the stimulus onset asynchrony of the auditory and visual stimulus (negative values indicate visual first). Solid lines: predictions by the model variant that assumes the uncertainty participants employed during perceptual inference did not match the actual uncertainty; model parameters are based on a joint model fit across all five tasks.

#### S3.2: Experiment 2 - Audiovisual spatial-discrimination task

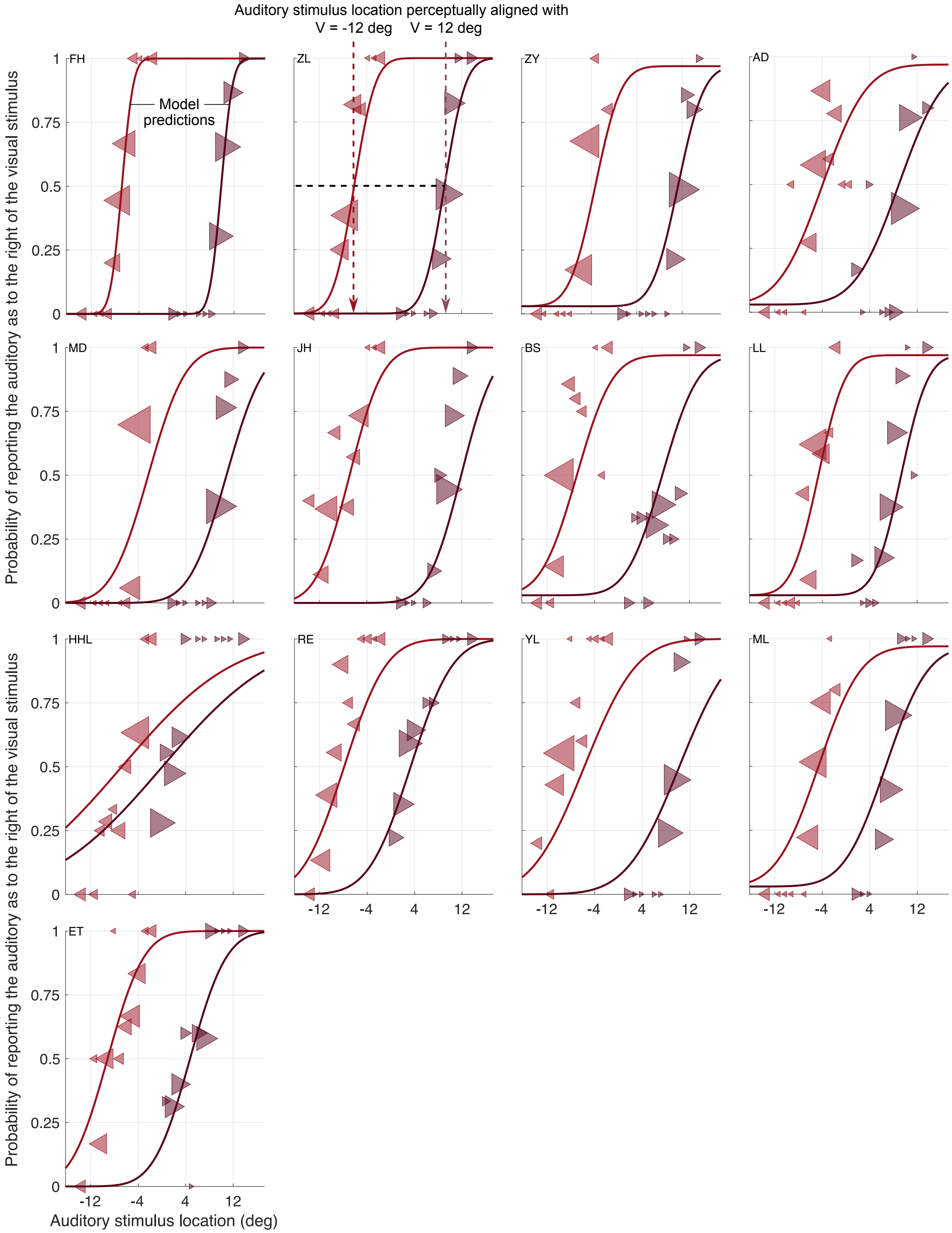


**Figure S5. Results of the audiovisual spatial-discrimination task (Expt. 2) for all participants.** Proportion of trials in which the auditory stimulus was reported as located to the right of the visual one as a function of the binned auditory stimulus location (markers, area proportional to the number of trials in each bin). Observed proportions and model predictions (lines) are shown separately for trials in which the visual stimulus was presented at -12 deg (leftward-pointing light red triangles) and trials in which the visual stimulus was presented at 12 deg (rightward-pointing dark red triangles). Curves show model predictions based on best-fitting parameters derived by jointly fitting each participants' data from all five tasks.

### S3.3: Experiment 3
#### S3.3.1: Localization practice

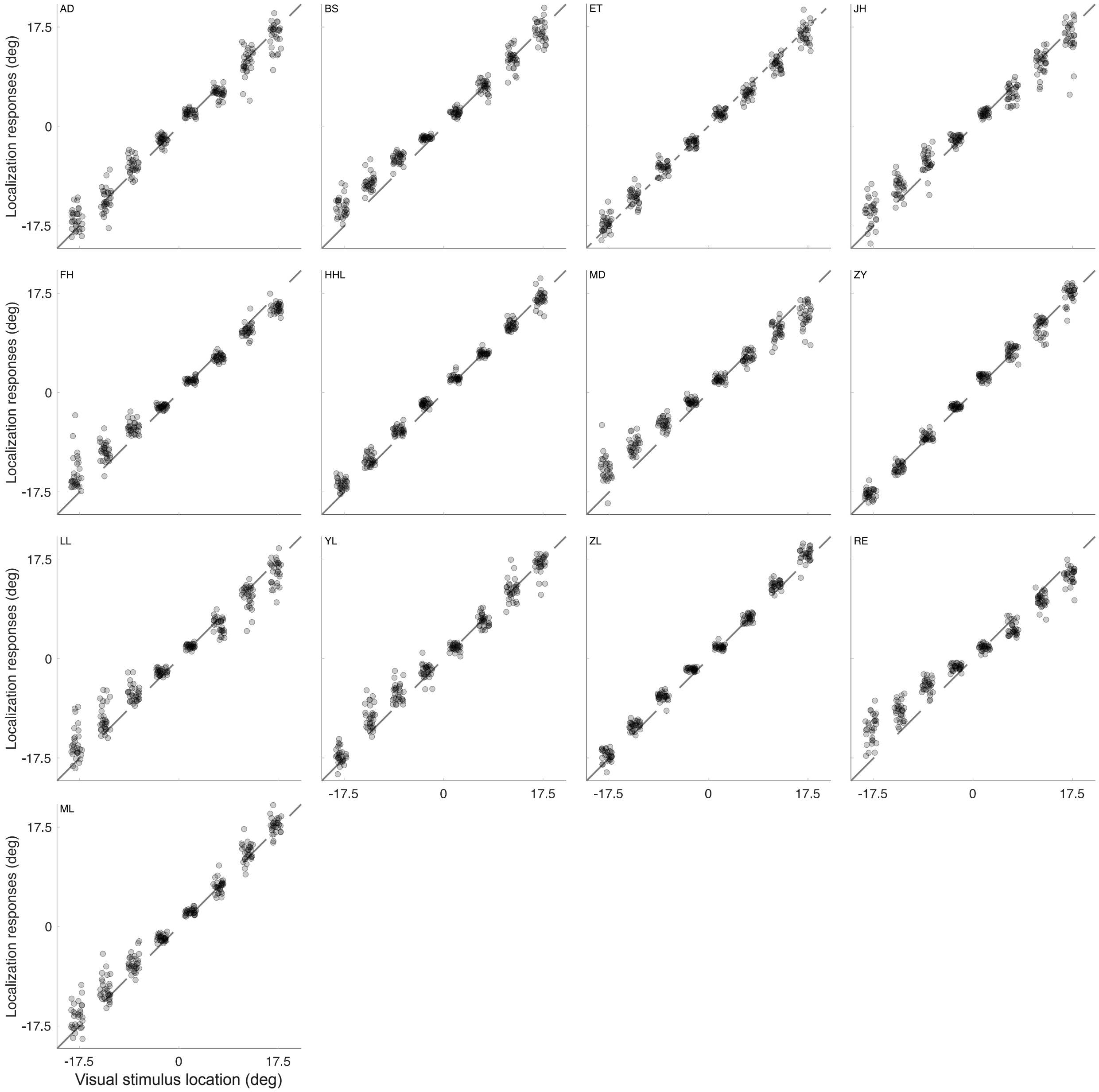


**Figure S6. Results of the localization-practice task for all participants.** Before completing the unimodal localization task of Experiment 3, participants practiced by localizing a small white square. Localization errors in this task were used to constrain the response-noise parameter. Dashed diagonal line: identity line. Data are slightly jittered horizontally for the purpose of visualization.

#### S3.3.2: Unimodal visual-localization task

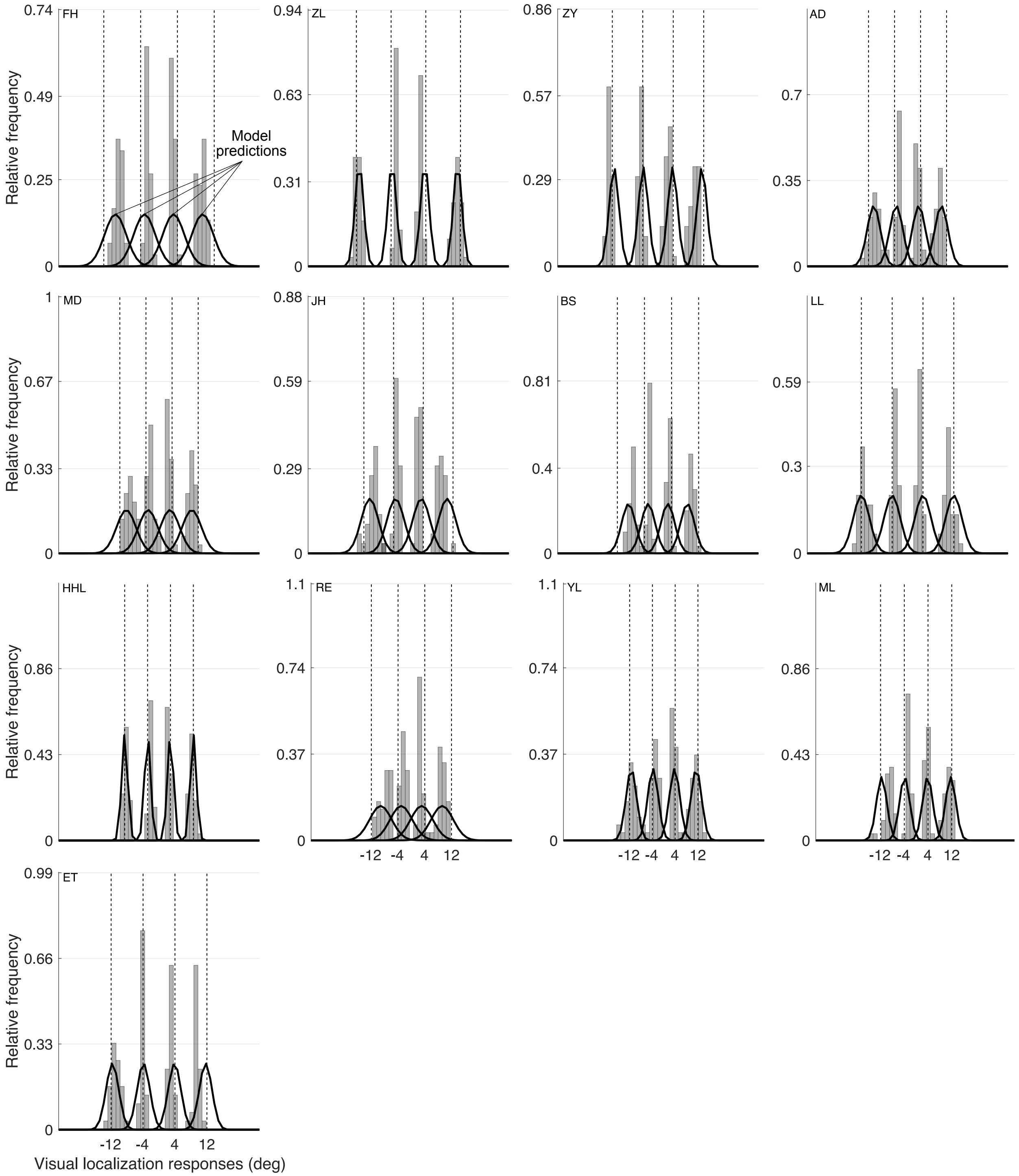


**Figure S7. Results of the unimodal visual-localization task for all participants.** Participants localized a visual stimulus (a Gaussian blob) presented at -12, -4, 4 or 12 deg relative to straight ahead. Solid lines: predictions by the model variant that assumes the uncertainty participants employed during perceptual inference did not match the actual uncertainty; model parameters are based on a joint model fit across all five tasks.

##### S3.3.3: Unimodal auditory-localization task

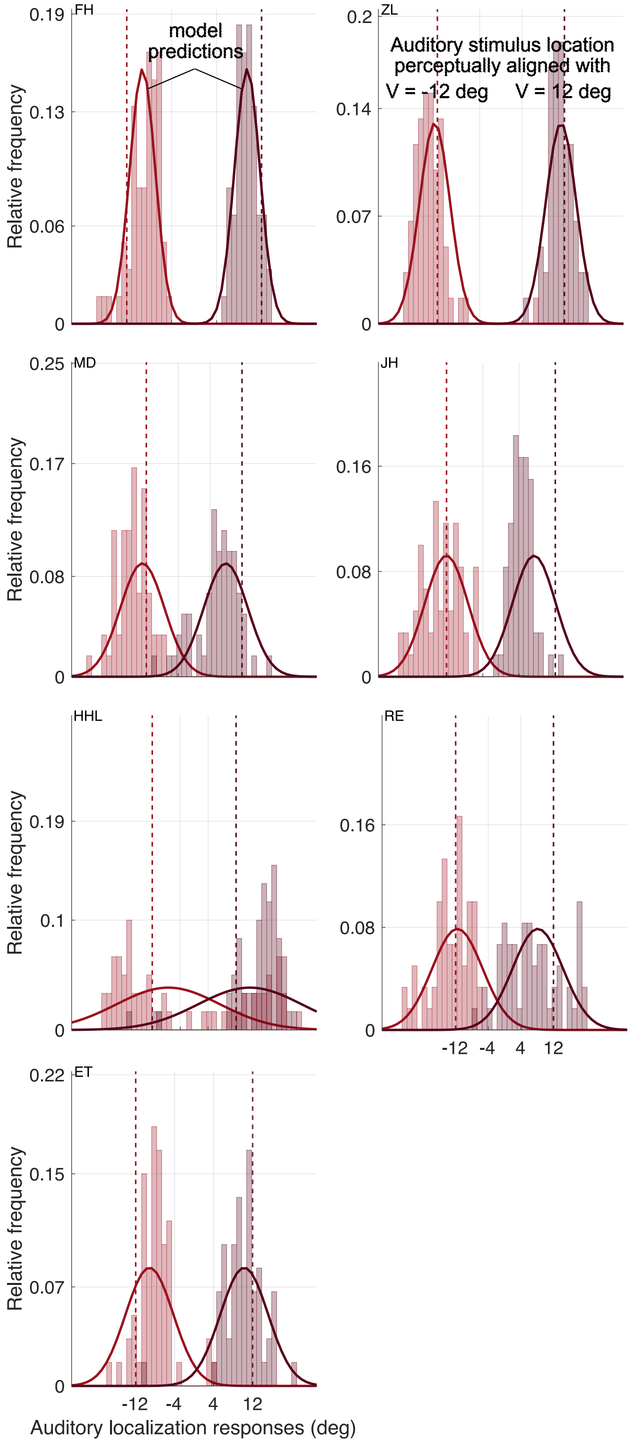


**Figure S8. Results of the unimodal auditory-localization task for all participants.** Participants localized auditory stimuli (100 ms-long noise bursts windowed using the positive half of a sine wave with a period of 200 ms) presented at one of two locations. These locations were identified for each participant to be perceptually aligned with a visual stimulus presented at -12 or 12 deg, respectively. *X*-axis labeling refers to this perceptual space. Solid lines: predictions of the model variant that assumes the uncertainty participants employed during perceptual inference did not match the actual uncertainty; model parameters are based on a joint model fit across all five tasks.

#### S3.4: Experiment 4
#### S3.4.1: Auditory-localization task in the bimodal context

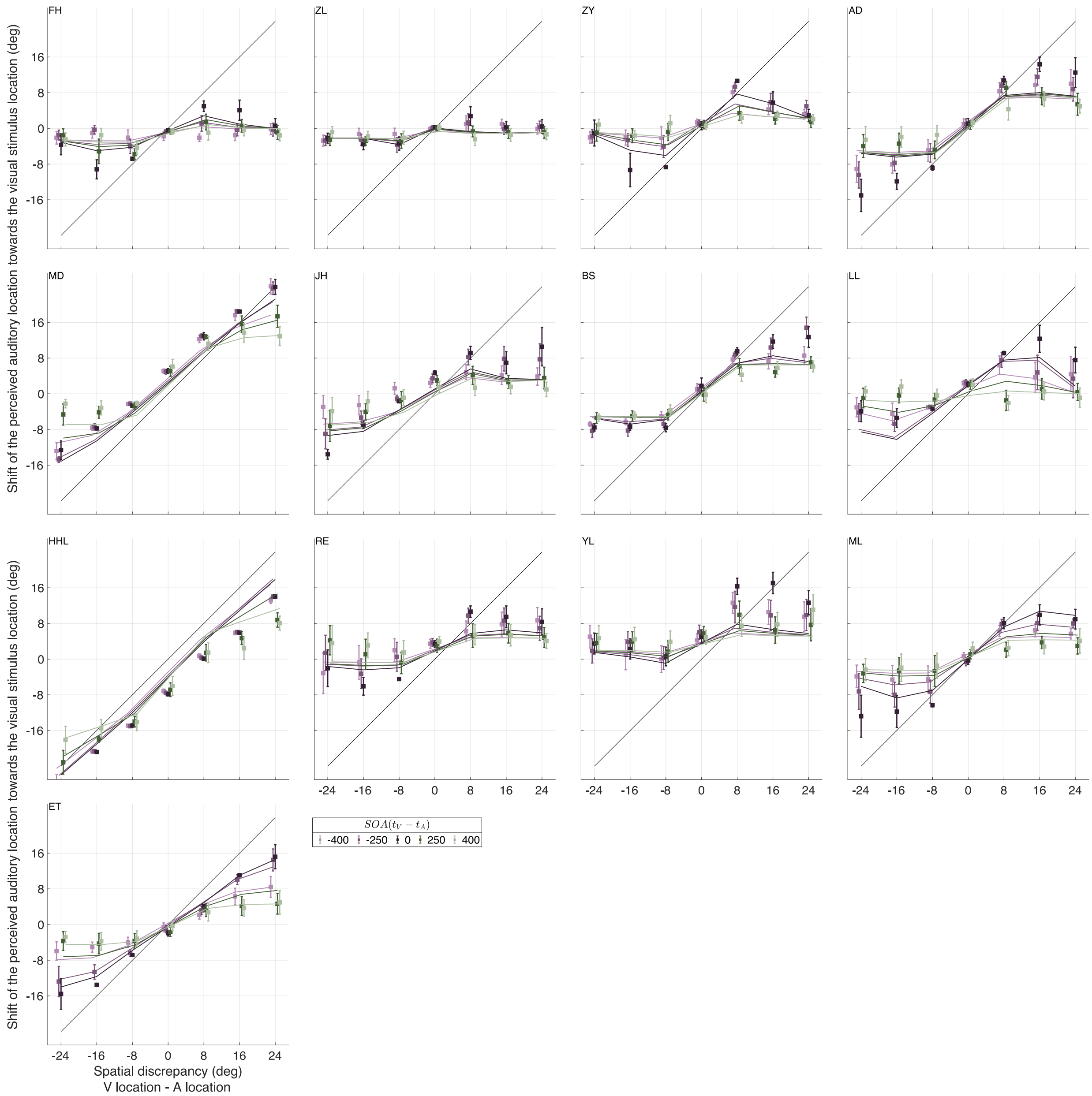


**Figure S9. Results of auditory spatial localization in audiovisual contexts for all participants.** Shifts of the perceived auditory location towards the visual stimulus location for each audiovisual temporal discrepancy as a function of audiovisual spatial discrepancy. Data are jittered slightly along the *x*-axis for legibility. Error bars: ±1 SD. Solid lines: predictions of the model variant that assumes the uncertainty participants employed during perceptual inference did not match the actual uncertainty; model parameters are based on a joint model fit across all five tasks.

Classical statistical analyses revealed that the absolute spatial and temporal discrepancies significantly affected performance in both the bimodal localization and the binary causal-inference judgments in Expt. 4. For the ventriloquism effect, the results showed that the absolute spatial and temporal discrepancies linearly ($\beta$ = -0.999, $t$ = -4.905, $p$ < 0.001) and quadratically ($\beta$ = 1.232, $t$ = 6.048, $p$ < 0.001) influenced participants' responses in a non-additive manner. For the binary causal-inference judgments, only a significant linear interaction of absolute spatial and temporal discrepancies emerged ($\beta$ = -0.263, $z$ = -2.871, $p$ = 0.004 , $p$ < 0.001).

| | Estimated $\beta$ (Confidence interval) | Test statistics and $p$-value |
|---|---|---|
| Intercept | 4.496 (3.118, 5.874) | $t = 6.626, p < 0.001$ *** |
| $\vert s_v - s_A \vert$ | 4.504 (4.105, 4.904) | $t = 22.114, p < 0.001$ *** |
| $\vert s_v - s_A \vert^2$ | -2.696 (-3.095, -2.297) | $t = -13.236, p < 0.001$ *** |
| $(t_v - t_A)$ | -0.888 (-1.002, -0.774) | $t = -15.256, p < 0.001$ *** |
| $(t_v - t_A)^2$ | -1.084 (-1.198, -0.970) | $t = -18.629, p < 0.001$ *** |
| $\vert s_v - s_A \vert : (t_v - t_A)$ | -0.999 (-1.398, -0.600) | $t = -4.905, p < 0.001$ *** |
| $\vert s_v - s_A \vert : (t_v - t_A)^2$ | -1.711 (-2.110, -1.312) | $t = -8.400, p < 0.001$ *** |
| $\vert s_v - s_A \vert^2 : (t_v - t_A)$ | 0.428 (0.029, 0.827) | $t = 2.103, p = 0.036$ * |
| $\vert s_v - s_A \vert^2 : (t_v - t_A)^2$ | 1.232 (0.833, 1.631) | $t = 6.048, p < 0.001$ *** |

**Table S1. Linear mixed-effects regression model (LMM) of ventriloquism effects (Fig. S9).** The temporal and absolute spatial discrepancy of the audiovisual stimulus pair were included as linear and quadratic predictors.

#### S3.4.2: Causal-inference judgments

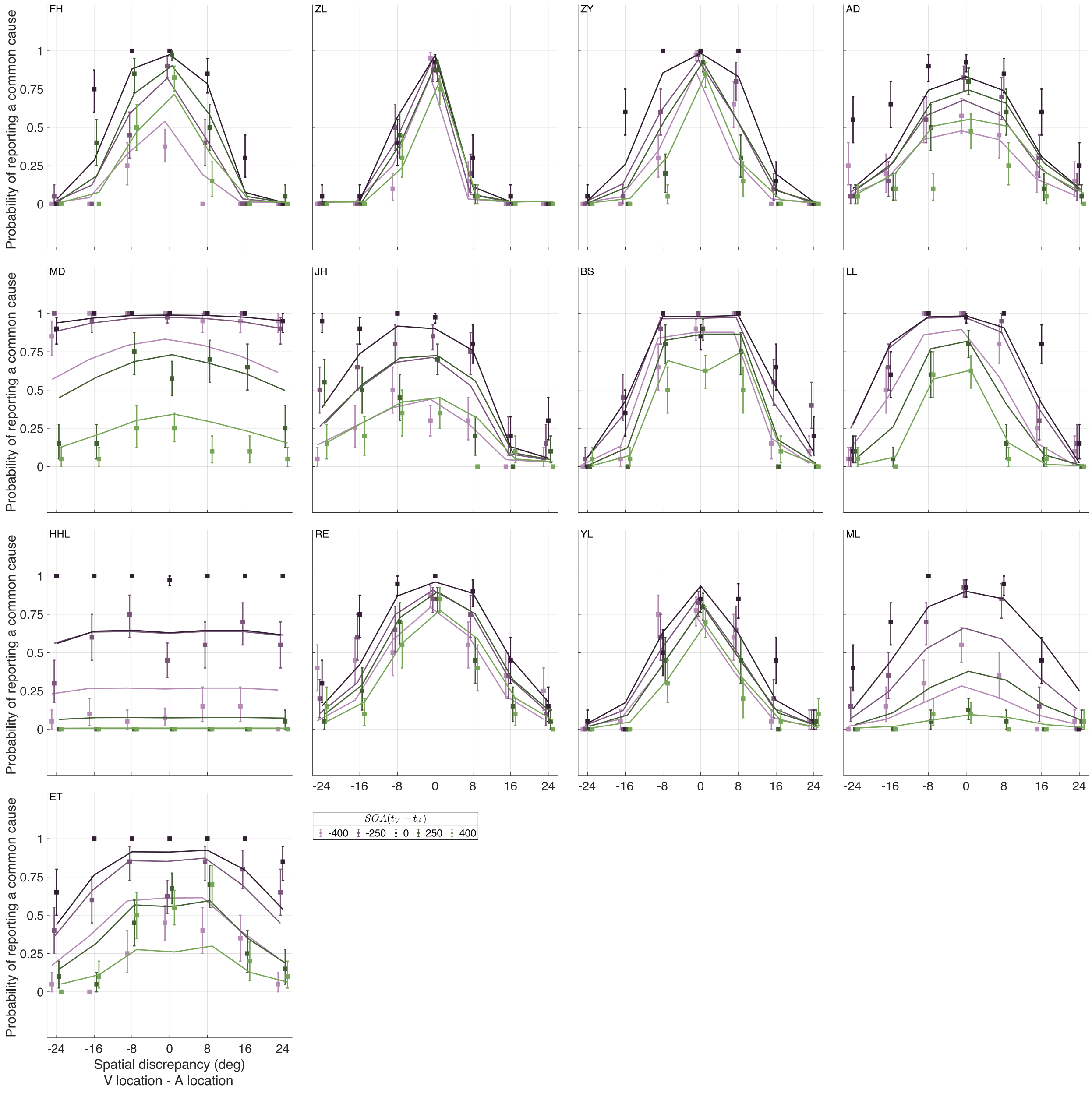


**Figure S10. Results of causal-inference judgments for all participants.** The probability of reporting a common cause for the visual and auditory stimulus as a function of their spatial discrepancy for each temporal discrepancy (colors). Error bars: ±1 SD. Solid lines: predictions of the model variant that assumes the uncertainty participants employed during perceptual inference did not match the actual uncertainty; model parameters are based on a joint model fit across all five tasks.

| | Estimated $\beta$ (Confidence interval) | Test statistics and $p$-value |
|---|---|---|
| Intercept | -0.519 (-0.948, -0.090) | $z$ = -2.553, $p$ = 0.011 * |
| $\lvert s_v - s_A \rvert$ | -1.513 (-1.696, -1.330) | $z$ = -16.205, $p$ < 0.001 *** |
| $\lvert s_v - s_A \rvert^2$ | 0.104 (-0.095, 0.301) | $z$ = 1.031, $p$ = 0.303 |
| $(t_v - t_A)$ | -0.622 (-0.687, -0.558) | $z$ = -18.929, $p$ < 0.001 *** |
| $(t_v - t_A)^2$ | -0.933 (-0.993, -0.874) | $z$ = -30.743, $p$ < 0.001 *** |
| $\lvert s_v - s_A \rvert : (t_v - t_A)$ | -0.263 (-0.442, -0.874) | $z$ = -2.871, $p$ = 0.004 ** |
| $\lvert s_v - s_A \rvert : (t_v - t_A)^2$ | -0.184 (-0.366, -0.002) | $z$ = -1.984, $p$ = 0.047 * |
| $\lvert s_v - s_A \rvert^2 : (t_v - t_A)$ | -0.076 (-0.131, 0.280) | $z$ = 0.721, $p$ = 0.471 |
| $\lvert s_v - s_A \rvert^2 : (t_v - t_A)^2$ | 0.115 (-0.078, 0.307) | $z$ = 1.168, $p$ = 0.243 |

**Table S2. Generalized linear mixed-effects regression model (GLMM) predicting explicit causal inference judgments in Experiment 4 (Fig. S10).** The temporal and absolute spatial discrepancy of the audiovisual stimulus pair were included as linear and quadratic predictors.

#### S4: Simulation results

Based on the best-fitting parameters, here we examine the effects of employing overconfident estimates of uncertainty on participants' performance. We simulated localization responses with the winning model (parameters specified in **Table S3**). In unimodal contexts, employing overconfident estimates of auditory spatial uncertainty leads to greater average accuracy, that is, lower mean localization error relative to the (remapped) location of the auditory stimulus (**Fig. S11A**). However, this improvement comes at the cost of precision, indicated by greater variability in location estimates across simulated trials (**Figs. S11B-C**). This tradeoff occurred also in bimodal contexts (**Figs. S11D-F**; see **Fig. S12** for the generalization of the effect across stimulus locations). Ultimately, employing overconfident estimates of uncertainty for perceptual

| Notation | Meaning | Value |
|---|---|---|
| $a_A$ | The proportional bias (location-dependent) in auditory spatial perception | 1 |
| $b_A$ | The constant bias (location-independent) in auditory spatial perception | 0 |
| $\sigma'_A$ | The sensory uncertainty of auditory-location measurements in unimodal trials | 5 deg |
| $\sigma'_{AV,A}$ | The sensory uncertainty of auditory-location measurements in bimodal trials | 16 deg |
| $\sigma'_V$ | The sensory uncertainty of visual-location measurements in unimodal and bimodal trials | 2 deg |
| $\mu'_P$ | The mean of the prior distribution (supra-modal) over stimulus location | 0 deg |
| $\sigma'_P$ | The width of the central prior (supra-modal) over stimulus location | 10 deg |
| $\sigma'_{\Delta_t}$ | The sensory uncertainty of stimulus-onset-asynchrony measurements | 158 ms |
| $b_{\Delta_t}$ | The constant bias in the measurements for estimating stimulus onset asynchrony | 0 ms |
| $\mu'_{P_{\Delta_t}}$ | The mean of the prior distribution over stimulus onset asynchrony given that the auditory and the visual stimuli came from one common source or two separate sources | 0 deg |
| $\sigma'_{P_{\Delta_t},C=1}$ | The width of the prior distribution over stimulus onset asynchrony for the common-cause scenario | 1 ms |
| $\sigma'_{P_{\Delta_t},C=2}$ | The width of the prior distribution over stimulus onset asynchrony for the separate-cause scenario | 800 ms |
| $p_{C=1}$ | Common-cause prior | 0.85 |
| $\epsilon_s$ | The internal criterion for making common-cause judgments in the spatial domain | 4 deg |
| $\epsilon_t$ | The internal criterion for making unity judgments in the temporal domain | 500 ms |

**Table S3. Parameter values for simulations based on the model that allows misestimation in auditory spatial uncertainty and audiovisual temporal uncertainty.**

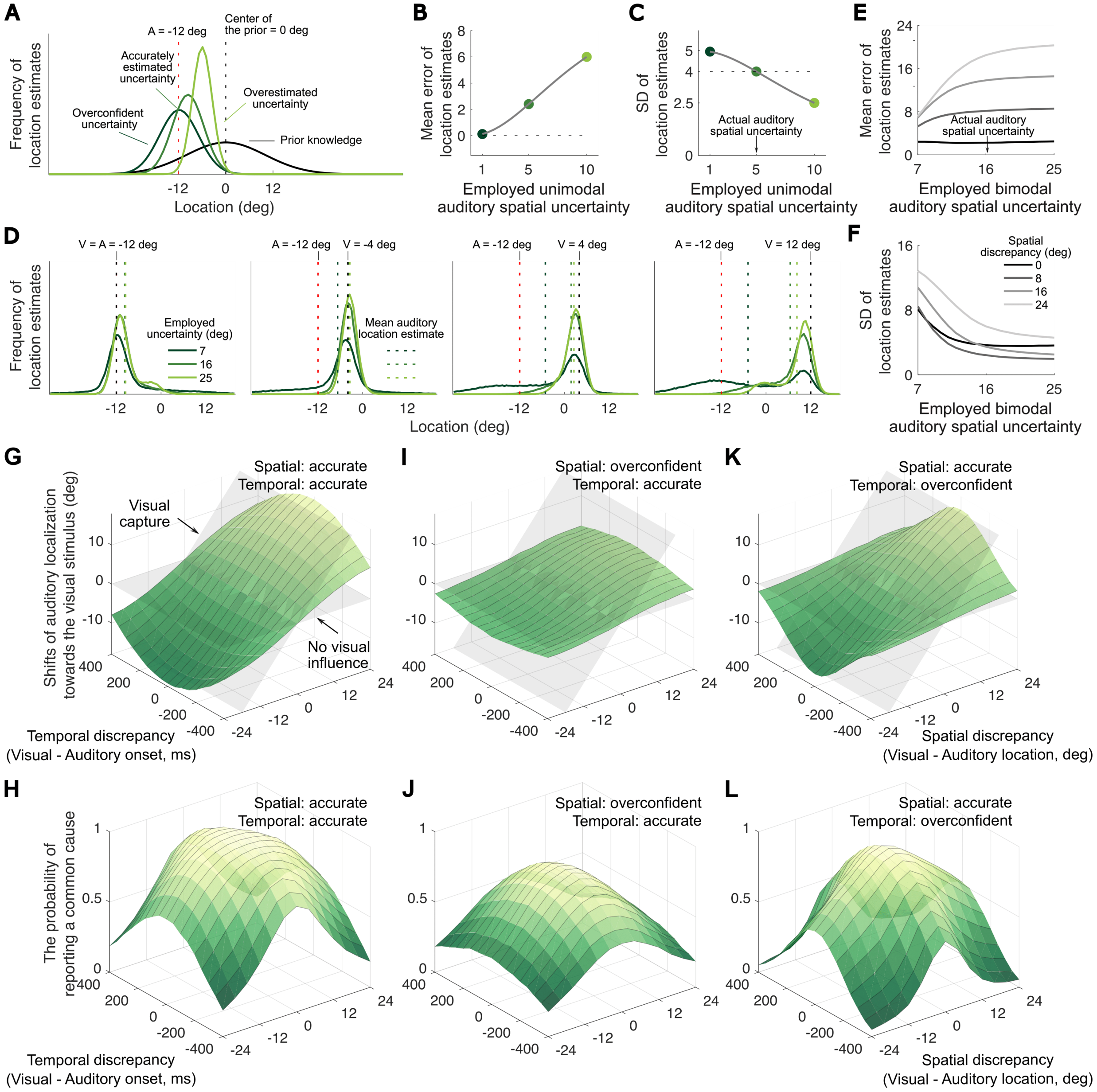


**Figure S11. Simulated effects of misestimated temporal and spatial auditory uncertainty on perceptual inference across the different tasks.** (A) Distributions of auditory-location estimates given an auditory stimulus presented alone at -12 deg, assuming the observer based the perceptual inference on an overconfident estimate of uncertainty (dark green), an accurate estimate of uncertainty (green) or overestimated their uncertainty (light green) associated with auditory spatial perception. For these simulations, the actual spatial uncertainty, the standard deviation of the measurement distribution, was set to 5 deg in the unimodal context. (B-C) Mean error and standard deviation of simulated auditory-location estimates as a function of the employed auditory spatial uncertainty, the standard deviation of the likelihood function. (D) Probability distributions of auditory-location estimates in bimodal contexts. Distributions were approximated using 100,000 Monte Carlo simulations. Auditory stimulus: -12 deg (red dashed line); visual stimulus: -12, -4, 4, 12 deg (black dashed line); temporal discrepancy (visual - auditory onset): 0 ms. (E-F) Mean error and standard deviation of auditory location estimates as a function of the employed spatial uncertainty, separated by the spatial discrepancy of the audiovisual stimulus pair (shades of grey). The actual spatial uncertainty was set to 16 deg for simulations of the bimodal context. The actual and employed audiovisual temporal uncertainty were set to be 158 and 75 ms, respectively. (G-L) Simulated shifts in auditory localization responses towards the accompanying visual stimulus and probabilities of reporting a common cause as a function of spatial and temporal discrepancy, assuming the observer employs accurate or inaccurate estimates of spatial and temporal uncertainty.

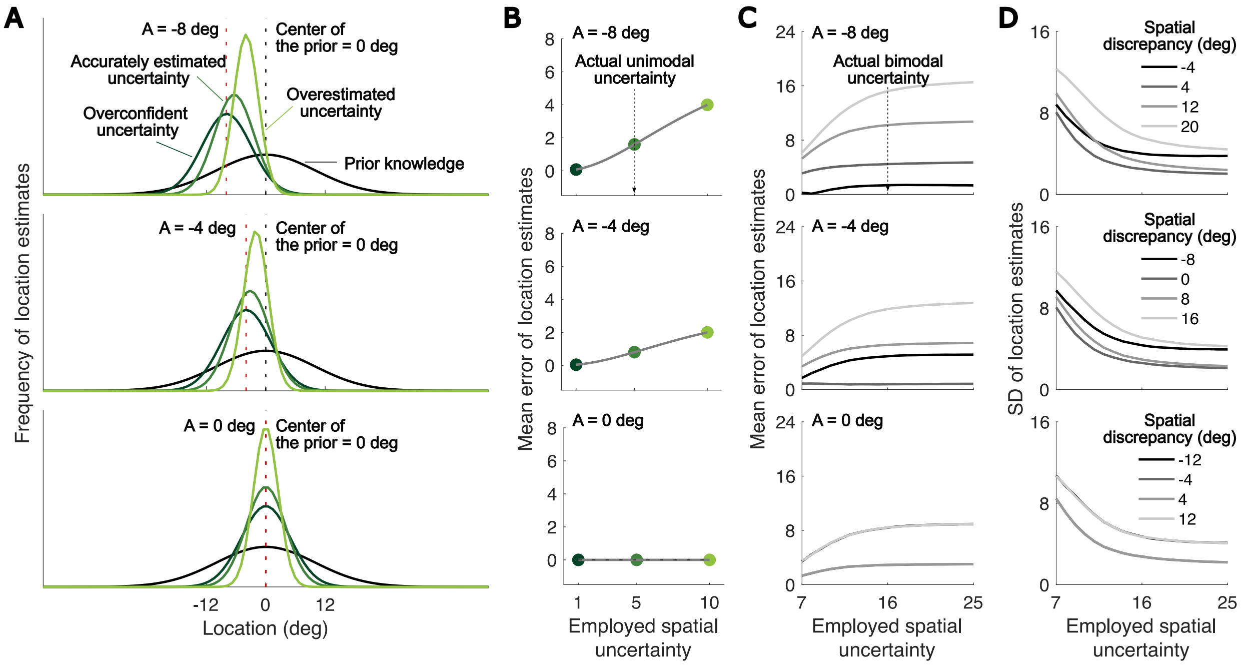


**Figure S12. Simulated effects of misestimated auditory spatial uncertainty on perceptual inferences in the unimodal and bimodal contexts.** (A) The distributions of auditory location estimates given the auditory stimulus presented alone at -8 deg (top panel), -4 deg (middle panel), and 0 deg (bottom panel) relative to straight ahead. The actual auditory spatial uncertainty was set to 5 deg in the unimodal contexts, while the employed uncertainty was varied to be either greater (light green), equivalent (green) or smaller (dark green) than the actual value. (B) The mean (absolute) error of auditory location estimates as a function of varying employed uncertainty given the auditory stimulus presented at three different locations (panels). The standard deviation of auditory location estimates remained unchanged across those auditory stimulus locations (see **Fig. S11C**). (C-D) The mean (absolute) error and the standard deviation of auditory location estimates as a function of the employed uncertainty, with different spatial discrepancy between the auditory stimulus pair (curves with different shades of grey), given those auditory stimulus locations (panels). The actual spatial uncertainty was set to 16 deg in the bimodal contexts. The actual and employed audiovisual temporal uncertainty were set to be 158 ms and 75 ms, respectively. Results were obtained using 10,000 Monte Carlo simulations.

inference leads to reduced audiovisual integration, manifested as smaller shifts of auditory location estimates towards the visual stimulus and a lower proportion of common-cause or unity percepts (**Figs. S11G-L**). The simulated reduction in audiovisual integration is driven more by overconfident estimates of auditory spatial than temporal uncertainty; although both estimates affect the posterior probability of the causal scenarios, auditory spatial uncertainty additionally influences auditory location estimates (see Section **S5**).

### S5: Modeling

#### S5.1: Formalization of the audiovisual temporal-order-judgment task (Experiment 1)

The audiovisual temporal-order judgment task was used to measure the actual temporal uncertainty associated with the perception of audiovisual stimulus pairs, $\sigma'_{\Delta_t}$, as well as observers' cross-modal temporal biases, $b_{\Delta_t}$.

In each trial, an auditory and a visual stimulus were presented with a stimulus onset asynchrony $s_{\Delta_t,i}$, where $i$ indexes the different SOAs ($i \in \{1,2,...,20\}$). Each presentation led to a noisy and biased measurement of the stimulus onset asynchrony, $m'_{\Delta_t}$, $m'_{\Delta_t} \sim \mathcal{N}(s'_{\Delta_t}, {\sigma'_{\Delta_t}}^2) = \mathcal{N}(s_{\Delta_t} + b_{\Delta_t}, {\sigma'_{\Delta_t}}^2)$. The spatial biases were assumed to vary across sessions, as they are greatly influenced by head positions, which differed slightly between sessions. Given the lack of a temporal reference point, the measurement of the temporal offset refers to the difference in visual relative to auditory arrival times ($t_V - t_A$), and we assumed Gaussian-distributed temporal noise in accordance with previous models (6).

We additionally assumed that in this task observers have a broad, uninformative prior over the temporal offsets between the stimuli. They had no reason to assume the auditory and visual stimulus would be presented at the same time or in a specific order as they were instructed to indicate the temporal order of the stimuli. As a consequence of this near-uniform prior, the posterior equals the likelihood and the observer's estimate of the temporal discrepancy in a single trial, $\hat{s}'_{\Delta_t}$, is identical to the measurement, $m'_{\Delta_t}$, and not impacted by the accuracy of the observer's estimate of audiovisual temporal uncertainty. The observer's decision in the temporal-order judgment task relies on the relative arrival times of the auditory and visual signals. Specifically, if $m'_{\Delta_t} > 0$, the observer infers that the auditory stimulus arrived first. Therefore, the probability of the observer inferring that the auditory stimulus preceded the visual stimulus, $p_A$, is

$$p_A = P(m'_{\Delta_t} > 0) = 1 - \Phi(0; s'_{\Delta_t}, {\sigma'_{\Delta_t}}^2) = 1 - \Phi(0; s_{\Delta_t} + b_{\Delta_t}, {\sigma'_{\Delta_t}}^2), \quad \text{(S1)}$$

where $\Phi(x; \mu, \sigma^2)$ refers to the cumulative distribution function of a Gaussian with mean $\mu$ and variance $\sigma^2$.

We additionally assume that the observer lapses at rate $\lambda_{TOJ}$. Therefore, the probability of reporting that the auditory stimulus occurred before the visual stimulus ($r = A$) for a given stimulus onset asynchrony $s_{\Delta_t}$ is equal to

$$p_{r=A} = 0.5\lambda_{TOJ} + (1 - \lambda_{TOJ})P(m'_{\Delta_t} > 0). \quad \text{(S2)}$$

#### S5.2: Formalization of the audiovisual spatial-discrimination task (Experiment 2)

The bimodal spatial-discrimination task was conducted to measure the actual spatial uncertainty associated with unimodally presented auditory and visual stimuli, $\sigma'_A$ and $\sigma'_V$, as well as the estimate of auditory uncertainty employed for perceptual inference, $\tilde{\sigma}'_A$. We assumed perceptual inference relied on an accurate estimate of visual spatial uncertainty, $\tilde{\sigma}'_V = \sigma'_V$, based on previous studies (7). Additionally, the task constrained estimates of other parameters that govern spatial perception, including biases in auditory spatial perception and a supra-modal prior over stimulus location.

In each trial, a visual standard stimulus, presented at one of two locations, $s_{V,n} \in \{-12°, 12°\}$, was paired with an auditory stimulus $s_{A,m}$, the location of which was determined by a staircase procedure. For each temporally separated stimulus pair, the model formalizes $p_{n,m}$, the probability of judging the auditory stimulus $s_{A,m}$ as to the right of the visual stimulus at location $s_{V,n}$ based on relation between the internal location estimates $\hat{s}'_{A,m}$ and $\hat{s}'_{V,n}$,

$$p_{n,m} = P(\hat{s}'_{A,m} > \hat{s}'_{V,n}) = P(\hat{s}'_{A,m} - \hat{s}'_{V,n} > 0). \quad \text{(S3)}$$

To further specify $p_{n,m}$, we have to derive the probability distribution of these two internal location estimates. We assume that each auditory stimulus leads to a noisy, biased internal measurement $m'_A$, $m'_A \sim \mathcal{N}(s'_A, {\sigma'_A}^2)$, where $s'_A = a_A s_A + b_A$, the remapped auditory stimulus location within an internal reference frame. Without loss of generality, we set $s'_V = s_V$, as visual and auditory biases trade off in our tasks.

The observer's estimate of the stimulus location relies on *a priori* knowledge (or assumptions). To avoid excessive model complexity, this spatial prior is assumed to be supra-modal, i.e., $s'_A \sim \mathcal{N}(\mu'_P, {\sigma'_P}^2)$ and $s'_V \sim \mathcal{N}(\mu'_P, {\sigma'_P}^2)$, and centered at perceived straight ahead, i.e., $\mu'_P = 0$. In every trial, the prior is combined with the likelihood, a Gaussian centered on the current measurement. The spread of the likelihood represents the observer's estimates of the stimulus' spatial uncertainty, $\tilde{\sigma}'_A$ and $\sigma_V$. The observer's location estimate, the mode of the posterior, is then a weighted combination of the mean of the prior, $\mu'_P$, and the measurement,

$$\begin{aligned}
\hat{s}'_{A,m} &= \frac{{\tilde{\sigma}'_A}^{-2}}{{\tilde{\sigma}'_A}^{-2} + {\sigma'_P}^{-2}} m'_{A,m} + \frac{{\sigma'_P}^{-2}}{{\tilde{\sigma}'_A}^{-2} + {\sigma'_P}^{-2}} \mu'_P, \text{ and} \\
\hat{s}'_{V,n} &= \frac{{\sigma'_V}^{-2}}{{\sigma'_V}^{-2} + {\sigma'_P}^{-2}} m'_{V,n} + \frac{{\sigma'_P}^{-2}}{{\sigma'_V}^{-2} + {\sigma'_P}^{-2}} \mu'_P .
\end{aligned} \quad \text{(S4)}$$

The probability distributions of the internal auditory and visual location estimates can then be written as

$$\hat{s}'_{A,m} \sim \mathcal{N}(\mu_{\hat{s}'_{A,m}}, \sigma^2_{\hat{s}'_A}) \text{ and } \hat{s}'_{V,n} \sim \mathcal{N}(\mu_{\hat{s}'_{V,n}}, \sigma^2_{\hat{s}'_V}), \tag{S5}$$

where

$$\begin{aligned}
\mu_{\hat{s}'_{A,m}} &= \frac{\tilde{\sigma}'^{-2}_A s'_{A,m} + \sigma'^{-2}_P \mu'_P}{\tilde{\sigma}'^{-2}_A + \sigma'^{-2}_P} = \frac{\tilde{\sigma}'^{-2}_A (a_A s_{A,m} + b_A) + \sigma'^{-2}_P \mu'_P}{\tilde{\sigma}'^{-2}_A + \sigma'^{-2}_P}, \\
\mu_{\hat{s}'_{V,n}} &= \frac{\tilde{\sigma}'^{-2}_V s'_{V,n} + \sigma'^{-2}_P \mu'_P}{\tilde{\sigma}'^{-2}_V + \sigma'^{-2}_P} = \frac{\tilde{\sigma}'^{-2}_V s_{V,n} + \sigma'^{-2}_P \mu'_P}{\tilde{\sigma}'^{-2}_V + \sigma'^{-2}_P}, \\
\sigma^2_{\hat{s}'_A} &= \sigma'^2_A \left( \frac{\tilde{\sigma}'^{-2}_A}{\tilde{\sigma}'^{-2}_A + \sigma'^{-2}_P} \right)^2 = \frac{\sigma'^2_A \sigma'^4_P}{(\tilde{\sigma}'^2_A + \sigma'^2_P)^2} \text{ and} \\
\sigma^2_{\hat{s}'_V} &= \sigma'^2_V \left( \frac{\sigma'^{-2}_V}{\sigma'^{-2}_V + \sigma'^{-2}_P} \right)^2 = \frac{\sigma'^2_V \sigma'^4_P}{(\sigma'^2_V + \sigma'^2_P)^2}.
\end{aligned} \tag{S6}$$

The probability distribution of the difference between the two location estimates $\hat{s}'_{A,m}$ and $\hat{s}'_{V,n}$ is

$$\hat{s}'_{A,m} - \hat{s}'_{V,n} \sim \mathcal{N}(\mu_{\hat{s}'_{A,m}} - \mu_{\hat{s}'_{V,n}}, \sigma^2_{\hat{s}'_A} + \sigma^2_{\hat{s}'_V}). \tag{S7}$$

Taken together, the probability of perceiving an auditory test stimulus at location $s_{A,m}$ to the right of a visual standard stimulus at location $s_{V,n}$ is

$$p_{n,m} = 1 - \Phi(0; \mu_{\hat{s}'_{A,m}} - \mu_{\hat{s}'_{V,n}}, \sigma^2_{\hat{s}'_A} + \sigma^2_{\hat{s}'_V}). \tag{S8}$$

We assume the observer lapses at rate $\lambda_{AV}$. Therefore, the probability of reporting an auditory stimulus at $s_{A,m}$ as located to the right of a visual stimulus at $s_{V,n}$ ($r_{n,m} = 1$) is equal to

$$p_{r_{n,m}=1} = 0.5\lambda_{AV} + (1 - \lambda_{AV})p_{n,m}. \tag{S9}$$

#### S5.3: Formalization of the unimodal auditory and visual spatial-localization tasks (Experiment 3)

The unimodal localization tasks were used to measure the actual spatial uncertainty associated with auditory and visual stimuli presented unimodally, $\sigma'_A$ and $\sigma'_V$, as well as the estimate of auditory spatial uncertainty used for perceptual inference, $\tilde{\sigma}'_A$. In addition, data from this task constrained biases in auditory spatial perception and the prior over stimulus locations. That is, all parameters constrained by the audiovisual spatial-discrimination task were also constrained by this task.

In this task, visual stimuli were presented at one of four locations in physical space ($s_{V,n} \in \{-12°, -4, 4, 12°\}$, where $n$ indexes visual location). Auditory stimuli were presented at one of two locations ($s_{A,m} \in \{s_{A,1}, s_{A,2}\}$, where $m$ indexes auditory location) selected for each participant to be perceptually aligned with the leftmost and rightmost visual locations (i.e., $s_{V,1} = -12^{\circ}$ and $s_{V,4} = 12^{\circ}$).

For each localization response, the observer adjusted a visual cursor until its location matched that of the estimated location of the stimulus, $\hat{s}'_{A,m}$ and $\hat{s}'_{V,n}$. We assumed that these localization responses are corrupted by perception-unrelated, Gaussian-distributed noise, $\sigma_r$, that arises due to the cognitive demand to hold the stimulus location in working memory and the sensorimotor requirements of adjusting a visual cursor. We assumed the response noise is unbiased, location-independent, and independent of the perceptual noise, that is,

$$r_{A,m} \sim \mathcal{N}(\mu_{\hat{s}'_{A,m}}, \sigma^2_{\hat{s}'_A} + \sigma^2_r) \text{ and } r_{V,n} \sim \mathcal{N}(\mu_{\hat{s}'_{V,n}}, \sigma^2_{\hat{s}'_V} + \sigma^2_r), \tag{S10}$$

where the terms $\mu_{\hat{s}'_A}$, $\mu_{\hat{s}'_V}$, $\sigma^2_{\hat{s}'_A}$ and $\sigma^2_{\hat{s}'_V}$ are defined in **Eq. S6**.

Preceding the regular spatial-localization task, participants familiarized themselves with the task by adjusting a visual cursor to match the location of a small, high contrast visual stimulus. Responses from this localization practice, $r_{V,o}$ where $o$ indexes the visual stimulus location, $s_{V,o}$, ($o \in \{1,2,...,8\}$), were used to estimate $\sigma^2_r$, as the visual stimuli presented during the practice should be associated with almost no spatial uncertainty:

$$r_{V,o} - s_{V,o} \sim \mathcal{N}(0,\sigma^2_r). \tag{S11}$$

#### S5.4: Formalization of the auditory spatial-localization task in a bimodal context (Experiment 4A)

This task was used to measure the spatial uncertainty associated with auditory, $\sigma'_{AV,A}$, and visual, $\sigma'_V$, stimuli presented as part of an audiovisual stimulus pair, the pair's temporal uncertainty, $\sigma'_{\Delta_t}$, and the estimates of auditory spatial, $\tilde{\sigma}'_{AV,A}$, and audiovisual temporal uncertainty, $\tilde{\sigma}'_{\Delta_t}$, that underlie perceptual inference. We assume that due to the attentional demands in this experiment's bimodal stimulus context, the auditory spatial uncertainty, $\sigma'_{AV,A}$, and its estimate, $\tilde{\sigma}'_{AV,A}$, might differ from the same parameters in unimodal contexts (8, 9). In contrast, given that the visual stimulus used in this study was highly reliable (see **Supplement S3.3.2**), the difference between the visual sensory noise for unimodal and bimodal presentations is assumed to be negligible. Therefore only one visual sensory uncertainty parameter, $\sigma'_V$, is implemented in our models. Additionally, the task constrained estimates of the other parameters that govern spatial and temporal perception in the tasks of Experiments 1, 2, and 3, including biases in auditory spatial and audiovisual temporal perception as well as the supra-modal prior over stimulus location.

In each trial, participants were presented with an audiovisual stimulus pair $s_{AV_l}$, where $l$ indexes different stimulus pairs used in the experiment. Each such stimulus pair consisted of an auditory component at location $s_{AV_l,A}$ and a visual component at location $s_{AV_l,V}$ with a temporal lag $s_{AV_l,\Delta_t}$ between the two components. The visual stimulus was presented at either one of the four locations, $s_{AV_l,V} \in \{-12°, -4°, 4°, 12°\}$; the auditory stimulus was presented at either one of the two locations, $s_{AV_l,A} \in \{s_{AV_l,A_1}, s_{AV_l,A_2}\}$, calculated to perceptually match $s_V = -12°$ and $s_V = 12°$, respectively, for each participant (**Supplement S2**). Using a scale of visual-equivalent locations, combining all the auditory and visual locations resulted in a total of seven spatial discrepancies ($[0°, \pm 8°, \pm 16°, \pm 24°]$). Auditory and visual stimuli were presented at one of five different stimulus-onset asynchronies $s_{AV_l,\Delta_t} \in \{-400, -250, 0, 250, 400\}$ ms. As a result, there were 40 unique audiovisual stimulus pairs $s_{AV_l}$ ($l \in \{1, 2, \ldots, 40\}$). The presentation of each such stimulus pair leads to three noisy sensory measurements, one of the auditory spatial location, $m'_{AV_l,A} \sim \mathcal{N}(s'_{AV_l,A}, {\sigma'_{AV,A}}^2) = \mathcal{N}(a_A s_{AV_l,A} + b_A, {\sigma'_{AV,A}}^2)$, one of the visual spatial location, $m'_{AV_l,V} \sim \mathcal{N}(s'_{AV_l,V}, {\sigma'_V}^2) = \mathcal{N}(s_{AV_l,V}, {\sigma'_V}^2)$, and one of the temporal discrepancy between the components, $m'_{AV_l,\Delta_t} \sim \mathcal{N}(s'_{AV_l,\Delta_t}, {\sigma'_{\Delta_t}}^2) = \mathcal{N}(s_{AV_l,\Delta_t} + b_{\Delta_t}, {\sigma'_{\Delta_t}}^2)$.

In this experiment the auditory and visual measurements might have either originated from two separate causes, $C = 2$, as in the other experiments, or from a common cause, $C = 1$. The posterior distributions over the range of possible auditory stimulus locations differ between these scenarios. In the common-cause scenario the auditory and visual stimuli are spatially aligned. Hence, the likelihood over visual stimulus locations provides information and should be integrated with the one over auditory stimulus locations and the supra-modal spatial prior. The location estimate of the auditory component $\hat{s}'_{AV_l,A,C=1}$ conditional on a common cause is thus:

$$\hat{s}'_{AV_l,A,C=1} = \frac{m'_{AV_l,A}{\tilde{\sigma}'_{AV,A}}^{-2} + m'_{AV_l,V}{\sigma'_V}^{-2} + \mu'_P{\sigma'_P}^{-2}}{{\tilde{\sigma}'_{AV,A}}^{-2} + {\sigma'_V}^{-2} + {\sigma'_P}^{-2}}. \quad \text{(S12)}$$

For the separate-causes scenario ($C = 2$), the posterior should only derive from the supra-modal prior and the auditory likelihood over space. The conditional location estimate of the auditory stimulus, $\hat{s}'_{AV_l,A,C=2}$, is:

$$\hat{s}'_{AV_l,A,C=2} = \frac{m'_{AV_l,A}{\tilde{\sigma}'_{AV,A}}^{-2} + \mu'_P{\sigma'_P}^{-2}}{{\tilde{\sigma}'_{AV,A}}^{-2} + {\sigma'_P}^{-2}} \quad \text{(S13)}$$

Note that we assumed that the observer uses the same (inaccurate) estimate of auditory spatial uncertainty for all perceptual inferences, i.e., to generate the conditional spatial estimates ($\hat{s}'_{AV,A,C=1}$, $\hat{s}'_{AV,A,C=2}$) and to derive the probability of a common cause.

The unconditional location estimate is derived by model averaging (10, 11). Specifically, the unconditional auditory location estimate $\hat{s}'_{AV_l,A}$ is the average of the two conditional auditory location estimates, the one given the common-cause scenario and the one given the separate-causes scenario, with each conditional estimate weighted by the posterior probability of the corresponding causal structure:

$$\hat{s}'_{AV_l,A} = \hat{s}'_{AV_l,A,C=1}P(C=1\,|\,m'_{AV_l,A}, m'_{AV_l,V}, m'_{AV_l,\Delta_t}) \\ +\hat{s}'_{AV_l,A,C=2}P(C=2\,|\,m'_{AV_l,A}, m'_{AV_l,V}, m'_{AV_l,\Delta_t})\,. \tag{S14}$$

The posterior probability of a common cause, $P(C=1\,|\,m'_{AV_l,A}, m'_{AV_l,A}, m'_{AV_l,\Delta_t})$, depends on the likelihood of a common cause given all three measurements as well as the common-cause prior $p_{C=1}$:

$$P(C=1\,|\,m'_{AV_l,A}, m'_{AV_l,V}, m'_{AV_l,\Delta_t}) = \\ \frac{P(m'_{AV_l,A}, m'_{AV_l,V}, m'_{AV_l,\Delta_t}\,|\,C=1)p_{C=1}}{P(m'_{AV_l,A}, m'_{AV_l,V}, m'_{AV_l,\Delta_t}\,|\,C=1)p_{C=1} + P(m'_{AV_l,A}, m'_{AV_l,V}, m'_{AV_l,\Delta_t}\,|\,C=2)(1-p_{C=1})}\,. \tag{S15}$$

The posterior probability of two separate sources is $1 - P(C=1\,|\,m'_{AV_l,A}, m'_{AV_l,V}, m'_{\Delta_t})$. Assuming independence of spatial and temporal sensory noise, the likelihood of a common cause (McGovern et al., 2016) equals

$$P(m'_{AV_l,A}, m'_{AV_l,V}, m'_{\Delta_t}\,|\,C=1) = P(m'_{AV_l,A}, m'_{AV_l,V}\,|\,C=1)P(m'_{AV_l,\Delta_t}\,|\,C=1) \\ = \left(\int P(m'_{AV_l,V}\,|\,s'_{AV_l})P(m'_{AV_l,A}\,|\,s'_{AV_l})P(s'_{AV_l})ds'_{AV_l}\right)\cdot\left(\int P(m'_{AV_l,\Delta_t}\,|\,s'_{\Delta_t})P(s'_{\Delta_t}\,|\,C=1)ds'_{\Delta_t}\right) \\ = \frac{1}{2\pi\sqrt{\sigma_V'^2\tilde{\sigma}_{AV,A}'^2 + \sigma_V'^2\sigma_P'^2 + \tilde{\sigma}_{AV,A}'^2\sigma_P'^2}} \\ \exp\left[-\frac{(m'_{AV_l,V}-m'_{AV_l,A})^2\sigma_P'^2 + (m'_{AV_l,V}-\mu'_P)^2\tilde{\sigma}_{AV,A}'^2 + (m'_{AV_l,A}-\mu'_P)^2\sigma_V'^2}{2(\sigma_V'^2\tilde{\sigma}_{AV,A}'^2 + \sigma_V'^2\sigma_P'^2 + \tilde{\sigma}_{AV,A}'^2\sigma_P'^2)}\right] \\ \cdot\frac{1}{\sqrt{2\pi(\tilde{\sigma}_{\Delta_t}'^2 + \sigma_{P_{\Delta_t},C=1}'^2)}}\exp\left[-\frac{(m'_{AV_l,\Delta_t}-\mu'_{P_{\Delta_t},C=1})^2}{2(\tilde{\sigma}_{\Delta_t}'^2 + \sigma_{P_{\Delta_t},C=1}'^2)}\right]. \tag{S16}$$

In the separate-causes scenario, all three measurements should be independent and thus

$$P(m'_{AV_l,A}, m'_{AV_l,V}, m'_{AV_l,\Delta_t}\,|\,C=2) = P(m'_{AV_l,A}\,|\,C=2)P(m'_{AV_l,V}\,|\,C=2)P(m'_{AV_l,\Delta_t}\,|\,C=2) \\ = \left(\int P(m'_{AV_l,V}\,|\,s'_V)P(s'_V)ds'_V\right)\left(\int P(m'_{AV_l,A}\,|\,s'_A)P(s'_A)ds'_A\right)\left(\int P(m'_{AV_l,\Delta_t}\,|\,s'_{\Delta_t})P(s'_{\Delta_t}\,|\,C=2)ds'_{\Delta_t}\right)$$

$$= \frac{1}{\sqrt{2\pi({\sigma'_V}^2 + {\sigma'_P}^2)}} \exp\left[-\frac{1}{2}\left(\frac{(m'_{AV_l,V} - \mu'_P)^2}{{\sigma'_V}^2 + {\sigma'_P}^2}\right)\right] \cdot \frac{1}{\sqrt{2\pi({\tilde{\sigma}'_{AV,A}}^2 + {\sigma'_P}^2)}} \exp\left[-\frac{1}{2}\left(\frac{(m'_{AV_l,A} - \mu'_P)^2}{{\tilde{\sigma}'_{AV,A}}^2 + {\sigma'_P}^2}\right)\right]$$

$$\cdot \frac{1}{\sqrt{2\pi({\tilde{\sigma}'_{\Delta_t}}^2 + {\sigma'_{P_{\Delta_t,C=2}}}^2)}} \exp\left[-\frac{(m'_{AV_l,\Delta_t} - \mu'_{P_{\Delta_t,C=2}})^2}{2({\tilde{\sigma}'_{\Delta_t}}^2 + {\sigma'_{P_{\Delta_t,C=2}}}^2)}\right]. \tag{S17}$$

When indicating the estimated location of the auditory stimulus, response noise might again corrupt the localization responses,

$$P(r_{AV_l,A} \mid m'_{AV_l,A}, m'_{AV_l,V}, m'_{\Delta_t}) = \phi(r_{AV_l,A}; \hat{s}'_{AV_l,A}, \sigma_r^2), \tag{S18}$$

where $r_{AV_l,A}$ is the indicated location, $\sigma_r$ the standard deviation of the perception-unrelated response noise, and $\phi$ is the Gaussian density function.

#### S5.5: Formalization of the causal-inference task (Experiment 4B)

This task was performed on the same stimuli as the auditory localization task in the bimodal context of Experiment 4. Directly after making their localization response, participants reported whether they perceived the audiovisual stimulus pair as having a common cause or separate causes. Hence, the task constrained the same parameters as described in the previous section: the spatial uncertainty associated with auditory, $\sigma'_{AV,A}$, and visual, $\sigma'_V$. stimuli presented as part of an audiovisual stimulus pair, the pair's temporal uncertainty, $\sigma'_{\Delta_t}$, and the estimates of auditory spatial, $\tilde{\sigma}'_{AV,A}$, and audiovisual temporal uncertainty, $\tilde{\sigma}'_{\Delta_t}$, used to make perceptual inferences. Additionally, the task constrained again estimates of the other parameters that govern spatial and temporal perception, including biases in auditory spatial and audiovisual temporal perception as well as a supra-modal prior over stimulus location. In addition, this task constrained scenario-dependent priors over audiovisual temporal offsets and internal decision thresholds.

Based on our previous results (9, 12), we assumed that participants adopted an estimate-based (heuristic) decision rule based on the distance between the auditory and visual location estimates, $\hat{s}'_{AV_l,A}$ and $\hat{s}'_{AV_l,V}$, as well as the estimate of the temporal offset, $\hat{s}'_{AV_l,\Delta_t}$. The visual location estimate is derived analogously to the auditory one (see **Eqs. S12-17**). The estimate of the temporal offset between the stimuli is based on model averaging just like the estimates of spatial location, i.e.,

$$\begin{aligned} \hat{s}'_{AV_l,\Delta_t} = \hat{s}'_{AV_l,\Delta_t,C=1} P(C=1 \mid m'_{AV_l,A}, m'_{AV_l,V}, m'_{AV_l,\Delta_t}) \\ + \hat{s}'_{AV_l,\Delta_t,C=2}\left(1 - P(C=1 \mid m'_{AV_l,A}, m'_{AV_l,V}, m'_{AV_l,\Delta_t})\right). \end{aligned} \tag{S19}$$

The two conditional estimates of the temporal offset again correspond to the modes of scenario-specific posterior distributions over the range of possible audiovisual offsets. However, unlike in the spatial domain, the prior over temporal offsets should differ with the causal scenario. Separate causes are associated with no prior information about the temporal offset between two unrelated events, implemented as a practically uninformative Gaussian prior with $\mu'_{P_{\Delta_t},C=2} = 0$ and $\sigma'_{P_{\Delta_t,C=2}} = 800$. In contrast, an auditory and a visual measurement that share a common cause should occur at the same time, i.e., with no or very little temporal offset.

Hence, we fixed $\mu'_{P_{\Delta_t},C=1} = 0$. Yet, the degree to which an observer would a priori assume any temporal offset in the common cause scenario might be specific to the individual (13), thus, we implemented $\sigma'_{P_{\Delta_t,C=1}}$ as a free parameter. The likelihood in either scenario is the same. The likelihood is centered on the measured temporal offset, $m'_{AV_l,\Delta_t}$, and its spread reflects the observer's estimate of temporal uncertainty, $\tilde{\sigma}'_{\Delta_t}$. Given that all priors and likelihoods are Gaussians, the two conditional estimates equal

$$\hat{s}'_{AV_l,\Delta_t,C=1} = \frac{m'_{AV_l,\Delta_t}\tilde{\sigma}'^{-2}_{\Delta_t} + \mu'_{P_{\Delta_t},C=1}\sigma'^{-2}_{P_{\Delta_t},C=1}}{\tilde{\sigma}'^{-2}_{\Delta_t} + \sigma'^{-2}_{P_{\Delta_t},C=1}} \text{ and}$$

$$\hat{s}'_{AV_l,\Delta_t,C=2} = \frac{m'_{AV_l,\Delta_t}\tilde{\sigma}'^{-2}_{\Delta_t} + \mu'_{P_{\Delta_t},C=2}\sigma'^{-2}_{P_{\Delta_t},C=2}}{\tilde{\sigma}'^{-2}_{\Delta_t} + \sigma'^{-2}_{P_{\Delta_t},C=2}}. \tag{S20}$$

If the distance between location estimates was below an internal criterion $\epsilon_s$ and the estimate for temporal discrepancy was below another internal criterion $\epsilon_t$, then participants would infer a common cause ($C = 1$). Otherwise, they would infer separate causes ($C = 2$). That is, this inference is

$$I_{C=1,AV_l} = \begin{cases} 1, & \text{if } |\hat{s}'_{AV_l,A} - \hat{s}'_{AV_l,V}| < \epsilon_s \text{ and } |\hat{s}'_{AV_l,\Delta_t}| < \epsilon_t, \\ 0, & \text{otherwise.} \end{cases} \tag{S21}$$

However, due to occasional lapses, $\lambda_{\text{unity}}$, the observer might mistakenly report separate causes even when the intended response is a common cause, and vice versa. That is, the actual report of the causal inference, $r_{I,AV_l}$, satisfies

$$\begin{aligned} &P(r_{C=1,AV_l} = 1 \,|\, m'_{AV_l,A}, m'_{AV_l,V}, m'_{AV_l,\Delta_t}) \\ &= \begin{cases} 1 - \lambda_{\text{unity}}, & \text{if } |\hat{s}'_{AV_l,A} - \hat{s}'_{AV_l,V}| < \epsilon_s \text{ and } |\hat{s}'_{AV_l,\Delta_t}| < \epsilon_t, \\ \lambda_{\text{unity}}, & \text{otherwise.} \end{cases} \end{aligned} \tag{S22}$$

#### S5.6: Model variants

We tested four different variants of the models outlined above. The model variants either assumed that the observer employed an accurate estimate of their auditory spatial uncertainty for perceptual inference ($\tilde{\sigma}'_A = \sigma'_A$ and $\tilde{\sigma}'_{AV,A} = \sigma'_{AV,A}$), an accurate estimate of audiovisual temporal uncertainty ($\tilde{\sigma}'_{\Delta_t} = \sigma'_{\Delta_t}$), accurate estimates of all uncertainties, or that neither the employed auditory spatial nor audiovisual matched the actual uncertainty associated with the stimuli. The model variants were tested for each set of models $M$, i.e., the assumptions of the

| Notation | | Meaning | | Constrained by | | | |
|---|---|---|---|---|---|---|---|
| | | | | Expt. 1 | Expt. 2 | Expt. 3 | Expt. 4 |
| Model variant-dependent parameters | $\tilde{\sigma}'_A$ | The employed uncertainty for auditory-location measurements in unimodal trials | | | ✓ | ✓ | ✓ |
| | $\tilde{\sigma}'_{AV,A}$ | The employed uncertainty for auditory-location measurements in bimodal trials | | | | | ✓ |
| | $\tilde{\sigma}'_{\Delta_t}$ | The employed sensory uncertainty for stimulus-onset-asynchrony measurements | | | | | ✓ |
| Model parameters consistent across all model variants | $\sigma'_A$ | The sensory uncertainty of auditory-location measurements in unimodal trials | | | ✓ | ✓ | |
| | $\sigma'_V$ | The sensory uncertainty of visual-location measurements in unimodal and bimodal trials | | | ✓ | ✓ | ✓ |
| | $\sigma'_P$ | The width of the central prior (supra-modal) over stimulus location | | | ✓ | ✓ | ✓ |
| | $a_A^i$ | The session-dependent proportional bias (location-dependent) in auditory spatial perception | $a_A^1$ | | ✓ | ✓ | |
| | | | $a_A^2$ | | | | ✓ |
| | | | $a_A^3$ | | | | ✓ |
| | $b_A^i$ | The session-dependent constant bias (location-independent) in auditory spatial perception | $b_A^1$ | | ✓ | ✓ | |
| | | | $b_A^2$ | | | | ✓ |
| | | | $b_A^3$ | | | | ✓ |
| | $b_{\Delta_t}$ | The constant bias in the measurements for estimating stimulus onset asynchrony | | ✓ | | | ✓ |
| | $\sigma'_{\Delta_t}$ | The sensory uncertainty of stimulus-onset-asynchrony measurements | | ✓ | | | ✓ |
| | $\sigma'_{AV,A}$ | The sensory uncertainty of auditory-location measurements in bimodal trials | | | | | ✓ |
| | $\sigma'_{P_{\Delta_t},C=1}$ | The width of the prior distribution over stimulus onset asynchrony for the common-cause scenario | | | | | ✓ |
| | $p_{C=1}$ | Common-cause prior | | | | | ✓ |
| | $\lambda_{unity}$ | The lapse rate for unity judgments | | | | | ✓ |
| | $\epsilon_s$ | The internal criterion for making common-cause judgments in the spatial domain | | | | | ✓ |
| | $\epsilon_t$ | The internal criterion for making common-cause judgments in the temporal domain | | | | | ✓ |

**Table S4. Summary of free model parameters.** Model parameters are split into those parameters that varied with the tested model variant and parameters that were consistent across all model variants.

model variant, such as accurate estimates of temporal uncertainty, $\tilde{\sigma}'_{\Delta_t} = \sigma'_{\Delta_t}$, were applied to the models of all five tasks.

#### S5.7: Model likelihoods

For a given set of models $M$, we jointly fit an observer's responses in the temporal-order judgment task in Experiment 1 ($X_1$), the audiovisual spatial-discrimination task in Experiment 2 ($X_2$), as well as to the localization responses for unimodally presented visual and auditory stimuli in Experiment 3 ($X_3$), and the localization responses to auditory stimuli presented as part of a stimulus pair along with the causal-inference task in Experiment 4 ($X_4$). We used a maximum-likelihood procedure to fit all models. That is, a set of free model parameters $\Theta_M$ (**Table S4**) was chosen to maximize the joint log likelihood

$$\log P(X \mid M, \Theta_M) = \sum_{k=1}^{5} \log P(X_k \mid M, \Theta_{M,k}), \tag{S23}$$

where $X_1, X_2, X_3, X_4, X_5 \subset X$ and $\Theta_{M,1}, \Theta_{M,2}, \Theta_{M,3}, \Theta_{M,4}, \Theta_{M,5} \subset \Theta_M$.

To reduce the number of free parameters estimated simultaneously, we separately fitted parameters capturing response noise and lapse rates. The perception-unrelated, response noise in all localization tasks, $\sigma_r$, was estimated based on the responses in the localization practice administered before Experiment 3 (**S5.3; Table S5**). We additionally estimated the lapse rates, $\lambda_{TOJ}$ and $\lambda_{AV}$, for the temporal-order judgment (Experiment 1) and the audiovisual spatial-discrimination task (Experiment 2) by separately fitting psychometric curves to these data (**S5.1** and **S5.2**). Finally, some parameters were fixed either based on theoretical assumptions or due to trade offs with other 'nuisance' parameters (**Table S6**).

##### S5.7.1: Model likelihoods for the audiovisual temporal-order-judgment task (Experiment 1)

In this task, participants indicated whether the auditory stimulus was presented before, $r = 1$, or after the visual stimulus, $r = 0$, of the audiovisual stimulus pair. For each trial, the likelihood of Model $M$ and a candidate parameter set $\Theta_1$ given the response $r$ is

$$P(r \mid M, \Theta_1) = p_{r=1}{}^{r} \cdot (1 - p_{r=1})^{1-r}, \tag{S24}$$

where $p_{r=1}$ is defined in **Eq. S2**. Thus, the log-likelihood given the responses across all trials (a total of 25 trials were tested for each of the 20 stimulus onset asynchronies) is

$$\log P(X_1 \mid M, \Theta_1) = \sum_{i=1}^{20} \sum_{t=1}^{25} \left[ r_{i(t)} \cdot \log p_{r_i=1} + (1 - r_{i(t)}) \cdot \log \left( 1 - p_{r_i=1} \right) \right]. \tag{S25}$$

| Notation | Meaning | Constrained by |
|---|---|---|
| $\lambda_{TOJ}$ | The lapse rate for discriminating the temporal order of visual and auditory stimuli | Expt. 1 |
| $\lambda_{AV}$ | The lapse rate for discriminating the relative spatial location of visual and auditory stimuli | Expt. 2 |
| $\sigma_r$ | Perception-independent noise | Expt. 3 (Localization practice) |

**Table S5. A summary of parameters that were fitted separately and then fixed during the joint model fit.**

| Notation | Meaning | Value |
|---|---|---|
| $\mu'_P$ | The mean of the prior distribution (supra-modal) over stimulus location | 0 |
| $\mu'_{P_{\Delta_t},C=1}$ | The mean of the prior distribution over stimulus onset asynchrony given that the auditory and the visual stimuli came from one common source | 0 |
| $\mu'_{P_{\Delta_t},C=2}$ | The mean of the prior distribution over stimulus onset asynchrony given that the auditory and the visual stimuli came from two separate sources | 0 |
| $\sigma'_{P_{\Delta_t},C=2}$ | The width of the prior distribution over stimulus onset asynchrony given that the auditory and the visual stimuli came from two separate sources | 800 |

**Table S6. A summary of parameters with fixed values.**

$p_{r_i=1}$ depends on the actual audiovisual temporal uncertainty associated with the stimulus onset asynchrony, $\sigma'_{\Delta_t}$, the audiovisual temporal bias, $b_{\Delta_t}$, and a lapse rate, $\lambda_{TOJ}$. The set of free parameters constrained by the data of the audiovisual temporal order task is $\Theta_1 = \{b_{\Delta_t}, \sigma'_{\Delta_t}, \lambda_{TOJ}\}$.

#### S5.7.2: Model likelihoods for the audiovisual spatial-discrimination task (Experiment 2)

In this task, participants indicated whether an auditory test stimulus was located to the left, $r_{n,m} = 0$, or to the right of a visual standard stimulus, $r_{n,m} = 1$. For each trial, the likelihood of a model $M$ and a candidate parameter set $\Theta_2$ given the response $r_{n,m}$ is

$$P(r_{n,m} \,|\, M, \Theta_2) = {p_{r_{n,m}=1}}^{r_{n,m}} \cdot (1 - p_{r_{n,m}=1})^{1-r_{n,m}}, \tag{S26}$$

where $p_{r_{n,m}=1}$ is defined in **Eq. S9**. Thus, the log likelihood given the responses across all tested visual and auditory locations is

$$\log P(X_2 \,|\, M, \Theta_2) = \sum_{n=1}^{2} \sum_{m} \left[ r_{n,m} \cdot \log(p_{r_{n,m}=1}) + (1 - r_{n,m}) \cdot \log\left(1 - p_{r_{n,m}=1}\right) \right]. \tag{S27}$$

$p_{r_{n,m}=1}$ depends on the spatial uncertainty associated with the auditory, $\sigma'_A$, and visual stimulus, $\sigma'_V$, as well as the estimate of auditory spatial uncertainty the observer employed for perceptual inference, $\tilde{\sigma}'_A$. Additionally, $p_{r_{n,m}=1}$ depends on the location-dependent and constant biases in auditory spatial perception, $a_A$ and $b_A$, the width of the supra-modal prior over stimulus location, $\sigma'_P$, and the lapse rate, $\lambda_{AV}$. The set of free parameters that were constrained by data from this task was $\Theta_2 = \{a_A^1, b_A^1, \tilde{\sigma}'_A, \sigma'_A, \sigma'_V, \sigma'_P, \lambda_{AV}\}$ for model variants that assume a mismatch between the actual and employed auditory spatial uncertainty, which is in turn reduced to $\Theta_2 = \{a_A^1, b_A^1, \sigma'_A, \sigma'_V, \sigma'_P, \lambda_{AV}\}$ for model variants that assume otherwise. Note that spatial biases were assumed to vary across sessions; since participants completed this experiment on the first session, the parameters being constrained are $a_A^1$ and $b_A^1$ (**Table S4**).

#### S5.7.3: Model likelihoods for the unimodal auditory and visual spatial-localization task

In each trial, participants localized either a unimodal visual or auditory stimulus, resulting in a cursor location setting $r_{A,m(t_A)}$ or $r_{V,n(t_V)}$. The localization responses are Gaussian-distributed (**Eq. S10**). From these distributions, we can compute the likelihood of a model $M$ and a candidate parameter set $\Theta_3$ from the Gaussian density functions evaluated at the observed localization responses $r_{A,m(t_A)}$ or $r_{V,n(t_V)}$:

$$P(r_{A,m(t_A)} \mid M, \Theta_3) = \phi\left(r_{A,m(t_A)}; \mu_{\hat{s}'_{A,m(t_A)}}, \sigma^2_{\hat{s}'_A} + \sigma_r^2\right), \text{ and}$$
$$P(r_{V,n(t_V)} \mid M, \Theta_3) = \phi\left(r_{V,n(t_V)}; \mu_{\hat{s}'_{V,n(t_V)}}, \sigma^2_{\hat{s}'_V} + \sigma_r^2\right). \quad \text{(S28)}$$

where $\mu_{\hat{s}'_A}$ and $\mu_{\hat{s}'_V}$ are defined in **Eq. S6**. The log-likelihood is the sum of the log-likelihoods across trials:

$$\log P(X_3 \mid M, \Theta_3) = \sum_{j=1}^{2} \sum_{t_A=1}^{40} \log P(r_{A,j(t_A)} \mid M, \Theta_3) + \sum_{k=1}^{4} \sum_{t_V=1}^{20} \log P(r_{V,k(t_V)} \mid M, \Theta_3). \quad \text{(S29)}$$

The log-likelihood depends on $\mu'_{\hat{s}_A}$ and $\mu'_{\hat{s}_V}$, which in turn depend on the actual auditory, $\sigma'_A$, and visual, $\sigma'_V$, spatial uncertainty during unimodal presentation, the estimate of auditory spatial uncertainty employed for perceptual inference, $\tilde{\sigma}'_A$, the spatial bias parameters, $a_A$ and $b_A$, as well as the width of the supra-modal prior over stimulus location, $\sigma'_P$. Therefore, the set of parameters constrained by localization responses in this task is $\Theta_3 = \{a_A^1, b_A^1, \tilde{\sigma}'_A, \sigma'_A, \sigma'_V, \sigma'_P, \sigma_r\}$ for model variants that assume a mismatch between the actual and employed auditory spatial uncertainty, which is in turn reduced to $\Theta_3 = \{a_A^1, b_A^1, \sigma'_A, \sigma'_V, \sigma'_P\}$ for model variants that assume otherwise. Note that spatial biases were assumed to vary across sessions; since participants completed both the audiovisual spatial discrimination experiment and this one on the first session, the parameters being constrained are $a_A^1$ and $b_A^1$ (**Table S4**).

**Localization task practice trials**. For each trial, the likelihood of the model parameters given a visual stimulus at location $s_{V,o(t)}$ and a subsequent response $r_{V,o(t)}$ is $\phi(r_{V,o(t)}; s_{V,o(t)}, \sigma_r)$. The only free parameter that was constrained by this task is $\sigma_r$. The maximum-likelihood estimate of $\sigma_r$ is

$$\sigma_r = \sqrt{\frac{\sum_{t=1}^{T} (r_{V,o(t)} - s_{V,o(t)})^2}{T}}. \tag{S30}$$

Outlier trials, identified as above or below 3 standard deviations of the mean, were excluded from this calculation. We estimated $\sigma_r$ and then fixed it for the joint model fit.

##### S5.7.4: Model likelihoods for the bimodal spatial-localization and causal-inference tasks

The main experiment consisted of 40 different audiovisual stimulus pairs, each of which was repeated 20 times ($t \in \{1, 2, \ldots, 20\}$). In each trial, after participants were presented with one of these stimulus pairs, they first localized the auditory component, which resulted in a cursor-location setting, $r_{AV_{l,t},A}$. Participants then made a causal-inference judgment, which resulted in a binary response $r_{C=1,AV_{l,t}}$ (1: common cause; 0: separate causes). The overall log-likelihood of a given model variant $M$ and set of model parameters $\Theta_{M,4}$ is the log-likelihood summed over all localization responses and causal-inference judgments:

$$\begin{aligned} &\log P(X_4 \mid M, \Theta_{M,4}) \\ &= \sum_{l=1}^{40} \sum_{t=1}^{20} \log\Big( \iiint P(r_{AV_{l,t},A}, r_{C=1,AV_{l,t}} \mid m'_{AV_l,A}, m'_{AV_l,V}, m'_{AV_l,\Delta_t}) \\ &\quad P(m'_{AV_l,A}, m'_{AV_l,V}, m'_{AV_l,\Delta_t} \mid s'_{AV_l,A}, s'_{AV_l,V}, s'_{AV_l,\Delta_t}) dm'_{AV_l,A} dm'_{AV_l,V} dm'_{AV_l,\Delta_t} \Big). \end{aligned} \tag{S31}$$

The three measurements, $m'_{AV_l,A}$, $m'_{AV_l,V}$ and $m'_{AV_l,\Delta_t}$, given the stimulus $s_{AV_l}$ are assumed to be independent, and thus the joint probability $P(m'_{AV_l,A}, m'_{AV_l,V}, m'_{AV_l,\Delta_t} \mid s'_{AV_l,A}, s'_{AV_l,V}, s'_{AV_l,\Delta_t})$ can be rewritten as the product of $P(m'_{AV_l,A} \mid s'_{AV_l,A})$, $P(m'_{AV_l,V} \mid s'_{AV_l,V})$ and $P(m'_{AV_l,\Delta_t} \mid s'_{AV_l,\Delta_t})$, and **Eq. S31** becomes

$$\begin{aligned} &\log P(X_4 \mid M, \Theta_{M,4}) \\ &= \sum_{l=1}^{40} \sum_{t=1}^{20} \log\Big( \iiint P(r_{AV_{l,t},A}, r_{C=1,AV_{l,t}} \mid m'_{AV_l,A}, m'_{AV_l,V}, m'_{AV_l,\Delta_t}) \\ &\quad P(m'_{AV_l,A} \mid s'_{AV_l,A}) P(m'_{AV_l,V} \mid s'_{AV_l,V}) P(m'_{AV_l,\Delta_t} \mid s'_{AV_l,\Delta_t}) dm'_{AV_l,A} dm'_{AV_l,V} dm'_{AV_l,\Delta_t} \Big). \end{aligned} \tag{S32}$$

Given that we assumed that all the measurements are Gaussian, the probability of landing on a specific measurement can be computed using the Gaussian density function, centered on the remapped stimulus location and using the actual sensory uncertainty, that is,

$$P(m'_{AV_l,A} \mid s'_{AV_l,A}) = \phi(m'_{AV_l,A}; a_A s_{AV_l,A} + b_A, {\sigma'_{AV,A}}^2),$$
$$P(m'_{AV_l,V} \mid s'_{AV_l,V}) = \phi(m'_{AV_l,V}; s_{AV_l,V}, {\sigma'_V}^2), \text{ and} \tag{S33}$$
$$P(m'_{AV_l,\Delta_t} \mid s'_{AV_l,\Delta_t}) = \phi(m'_{AV_l,\Delta_t}; s_{AV_l,\Delta_t} + b_{\Delta_t}, {\sigma'_{\Delta_t}}^2).$$

For all of our model variants, the measurements lead deterministically to both the auditory location estimate and explicit causal-inference response. The only probabilistic processes are the motor noise (for the localization responses) and lapses (for the explicit causal-inference judgments). Thus, **Eq. S32** can be rewritten as

$$\begin{aligned}
&\log P(X_4 \mid M, \Theta_{M,4}) \\
&= \sum_{l=1}^{40} \sum_{t=1}^{20} \log\Big( \iiint P(r_{AV_{l,t},A} \mid m'_{AV_l,A}, m'_{AV_l,V}, m'_{AV_l,\Delta_t}) P(r_{C=1,AV_{l,t}} \mid m'_{AV_l,A}, m'_{AV_l,V}, m'_{AV_l,\Delta_t}) \\
&\phi(m'_{AV_l,A}; a_A s_{AV_l,A} + b_A, {\sigma'_{AV,A}}^2) \phi(m'_{AV_l,V}; s_{AV_l,V}, {\sigma'_V}^2) \\
&\phi(m'_{AV_l,\Delta_t}; s_{AV_l,\Delta_t} + b_{\Delta_t}, {\sigma'_{\Delta_t}}^2) dm'_{AV_l,A} dm'_{AV_l,V} dm'_{AV_l,\Delta_t} \Big),
\end{aligned} \tag{S34}$$

where the term $P(r_{AV_{l,t},A} \mid m'_{AV_l,A}, m'_{AV_l,V}, m'_{AV_l,\Delta_t})$ is defined in **Eq. S18**, $P(r_{C=1,AV_{l,t}} \mid m'_{AV_l,A}, m'_{AV_l,V}, m'_{AV_l,\Delta_t})$ is defined in **Eq. S22**.

The log-likelihood depends on the auditory localization responses $r_{AV_{l,t},A}$ and causal-inference judgments $r_{C=l,AV_{l,t}}$, which in turn depend on the actual auditory, $\sigma'_{AV,A}$, and visual, $\sigma'_V$, spatial uncertainty and audiovisual temporal uncertainty, $\sigma'_{\Delta_t}$, as well as the estimates of auditory spatial, $\tilde{\sigma}'_{AV,A}$, and audiovisual temporal uncertainty, $\tilde{\sigma}'_{\Delta_t}$, employed for perceptual inference. Additionally, the auditory localization responses depend on spatial biases, $a_A$ and $b_A$, the temporal bias, $b_{\Delta_t}$, the width of the supra-modal prior over stimulus location, $\sigma'_P$, and the common-cause prior, $p_{C=1}$. The causal inference judgments depend additionally on the width of the prior over temporal discrepancy conditioned on the common-cause scenario, $\sigma'_{P_{\Delta_t},C=1}$, internal criteria for unity judgments ($\epsilon_s$ and $\epsilon_t$), and a lapse rate for unity judgments ($\lambda_{\text{unity}}$). Therefore, the set of parameters constrained by the localization responses and unity judgments is $\Theta_4 = \{a_A^2, a_A^3, b_A^2, b_A^3, b_{\Delta_t}, \tilde{\sigma}'_{AV,A}, \sigma'_{AV,A}, \sigma'_V, \tilde{\sigma}'_{\Delta_t}, \sigma'_{\Delta_t}, \sigma'_P, \sigma'_{P_{\Delta_t},C=1}, p_{C=1}, \epsilon_s, \epsilon_t, \lambda_{\text{unity}}\}$. The number of free parameters is reduced depending on whether the model variant $M$ assumes the employed auditory spatial uncertainties match the actual ones (i.e., $\tilde{\sigma}'_{AV,A} = \sigma'_{AV,A}$), the employed temporal uncertainty matches the actual one (i.e., $\tilde{\sigma}'_{\Delta_t} = \sigma'_{\Delta_t}$), or both (i.e., $\tilde{\sigma}'_{AV,A} = \sigma'_{AV,A}$ and $\tilde{\sigma}'_{\Delta_t} = \sigma'_{\Delta_t}$). Note that spatial biases were assumed to vary across sessions;

since participants completed 2/3 of this experiment on the second session and the rest 1/3 on the third session, the parameters being constrained are $a_A^2, a_A^3$ and $b_A^2, b_A^3$ (**Table S4**).

#### S5.8: Model fitting

##### S5.8.1: Parameter estimation

For each model $M$, we searched for the set of model parameters that minimized the joint negative log-likelihood using the BADS toolbox (14). To deal with parameter values that correspond to a local minimum of the negative log-likelihood, we ran BADS multiple times initiating the search with 20 different starting points, randomly chosen from a $D_M$-dimensional space, where $D_M$ is the number of free parameters for model $M$. The final set of parameter estimates is the set with the maximum joint likelihood across all runs of the fitting procedure.

##### S5.8.2: Model comparison

To compare model performance quantitatively, we computed the Akaike information criterion (AIC) (15) for each model variant, separately for each participant. The best-fitting model variant corresponds to the model variant with the minimal AIC value. We then computed relative scores, $\Delta_{AIC}$, which capture the fit of each model variant relative to the best-fitting one.

### S6: Model-estimated uncertainties and model comparison

#### S6.1: Model-estimated actual and employed uncertainties for perceptual inference

| | Actual audiovisual temporal uncertainty | Employed audiovisual temporal uncertainty | Actual auditory spatial uncertainty (unimodal) | Employed auditory spatial uncertainty (unimodal) | Actual auditory spatial uncertainty (bimodal) | Employed auditory spatial uncertainty (bimodal) |
|---|---|---|---|---|---|---|
| | $\sigma'_{\Delta t}$ | $\tilde{\sigma}'_{\Delta t}$ | $\sigma'_{A}$ | $\tilde{\sigma}'_{A}$ | $\sigma'_{AV,A}$ | $\tilde{\sigma}'_{AV,A}$ |
| FH | 182.062 | 78.528 | 2.755 | 3.902 | 11.452 | 5.927 |
| ZL | 80.476 | 81.772 | 3.585 | 1.247 | 3.925 | 1.476 |
| ZY | 167.624 | 18.359 | 3.454 | 0.245 | 7.387 | 6.244 |
| AD | 301.120 | 82.566 | 9.773 | 2.683 | 16.239 | 5.183 |
| MD | 147.993 | 81.602 | 5.106 | 0.745 | 6.495 | 13.218 |
| JH | 189.607 | 99.221 | 5.723 | 1.704 | 15.313 | 5.482 |
| BS | 64.556 | 81.657 | 4.680 | 0.317 | 7.513 | 3.078 |
| LL | 135.674 | 70.468 | 5.744 | 6.967 | 5.064 | 6.561 |
| HHL | 147.121 | 156.433 | 14.883 | 3.886 | 10.518 | **44.915** |
| RE | 147.075 | 99.583 | 5.422 | 0.113 | 14.562 | 4.271 |
| YL | 198.502 | 69.977 | 9.229 | 2.302 | 16.421 | 5.814 |
| ML | 144.541 | 55.990 | 5.478 | 0.115 | 13.981 | 8.538 |
| ET | 176.960 | 85.525 | 4.731 | 0.113 | 13.044 | 12.234 |

**Table S7. Model-estimated actual and employed uncertainties during perceptual inferences for each participant.** First column: participants' initials. Highlighted row: identified outlier.

#### S6.2: Model comparison

| | Model 1: $\tilde{\sigma}'_{\Delta t} = \sigma'_{\Delta t}$ $\tilde{\sigma}'_{A} = \sigma'_{A}$ $\tilde{\sigma}'_{AV,A} = \sigma'_{AV,A}$ | Model 2: $\tilde{\sigma}'_{\Delta t} = \sigma'_{\Delta t}$ $\tilde{\sigma}'_{A} \neq \sigma'_{A}$ $\tilde{\sigma}'_{AV,A} \neq \sigma'_{AV,A}$ | Model 3: $\tilde{\sigma}'_{\Delta t} \neq \sigma'_{\Delta t}$ $\tilde{\sigma}'_{A} = \sigma'_{A}$ $\tilde{\sigma}'_{AV,A} = \sigma'_{AV,A}$ | Model 4: $\tilde{\sigma}'_{\Delta t} \neq \sigma'_{\Delta t}$ $\tilde{\sigma}'_{A} \neq \sigma'_{A}$ $\tilde{\sigma}'_{AV,A} \neq \sigma'_{AV,A}$ |
|---|---|---|---|---|
| FH | 6135.4 | 6042.5 | 6020.5 | 6003.2 |
| ZL | 5266.9 | 5232.8 | 5269.4 | 5243.1 |
| ZY | 5983.5 | 5983.8 | 5920.6 | 5916.5 |
| AD | 7274.0 | 7112.1 | 7178.6 | 7098.8 |
| MD | 6249.4 | 6228.0 | 6250.7 | 6204.3 |
| JH | 6985.6 | 6928.3 | 6909.9 | 6836.3 |
| BS | 6124.4 | 5989.7 | 6126.0 | 5981.2 |
| LL | 6114.0 | 6117.3 | 6104.5 | 6108.4 |
| HHL | 6128.0 | 6054.4 | 6131.8 | 6059.8 |
| RE | 7218.8 | 7082.6 | 7156.5 | 7078.8 |
| YL | 6986.5 | 6975.2 | 6964.3 | 6952.4 |
| ML | 6833.8 | 6831.5 | 6789.8 | 6781.5 |
| ET | 6346.1 | 6316.1 | 6226.9 | 6223.7 |

**Table S8. AIC values of all model variants for each participant.** First column: participants' initials. Model 1 assumes participants employed accurate audiovisual temporal and auditory spatial uncertainties. Model 2 assumes participants employed accurate audiovisual temporal uncertainty but inaccurate auditory spatial uncertainty in both unimodal and bimodal contexts. Model 3 assumes participants employed inaccurate audiovisual temporal uncertainty but accurate auditory spatial uncertainty in both unimodal and bimodal contexts. Model 4 assumes participants employed inaccurate audiovisual temporal and auditory spatial uncertainties.

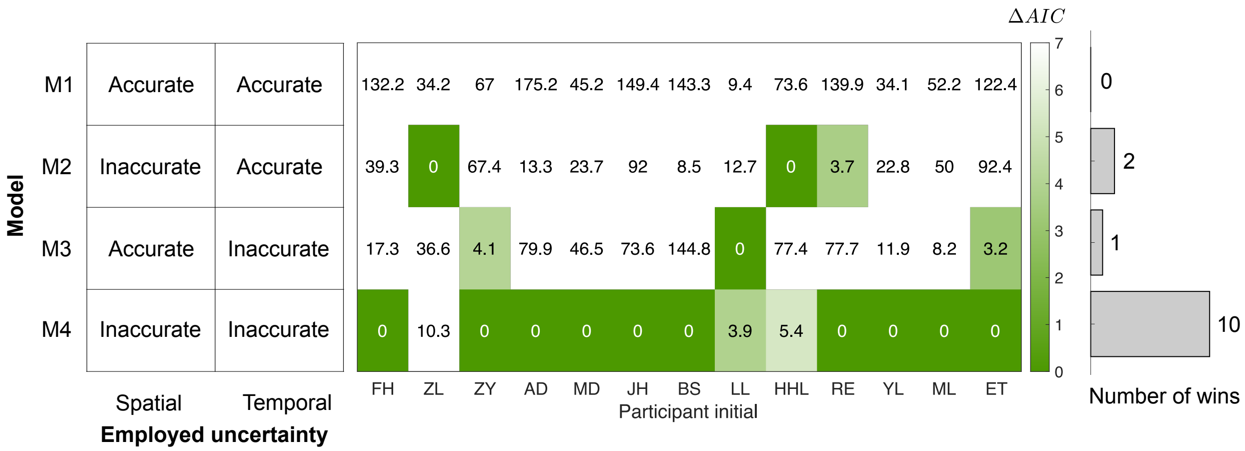


**Figure S13. Comparison between the four model variants for all participants.** Color key of the heat map: $\Delta_{AIC}$ (more saturated green represents better model performance). Color axis is clipped at 7 for the purpose of visualization.
